# Disorganized Inhibitory Dynamics in Hippocampal area CA1 of 22q11.2 Deletion Mutant Mice

**DOI:** 10.1101/2024.04.28.591464

**Authors:** Stephanie A. Herrlinger, Bovey Y Rao, Margaret E. Conde Paredes, Anna L. Tuttman, Haroon Arain, Erdem Varol, Joseph A. Gogos, Attila Losonczy

## Abstract

Individuals with the 22q11.2 deletion syndrome, one of the strongest genetic risk factors for schizophrenia, demonstrate cognitive impairments, including episodic memory dysfunction. Place cell activity of excitatory pyramidal neurons in the hippocampus supporting episodic memory is impaired in a mouse model for the 22q11.2 deletion (*Df(16)A^+/-^*). While excitatory dynamics are under tight inhibitory control by multiple subtypes of GABAergic interneurons, previous studies have predominantly focused on a single subtype of PV-expressing interneurons; there have not yet been studies describing the functional relationships between molecularly identified inhibitory types in *Df(16)A^+/-^*mice. Here, we examined interneuron subtype-specific activity dynamics in the dorsal hippocampal area CA1 of *Df(16)A^+/-^* mice during random foraging and spatial reward navigation tasks. Capitalizing on 3D acousto-optical deflector two-photon microscopy with *post hoc* immunohistochemical identification, we found that multiple interneuron types exhibit aberrant responses to reward locations and delayed reward enrichment extinction. *Df(16)A^+/-^* inhibitory interneurons also carry markedly reduced spatial information in a subtype-dependent manner. We observed task-dependent changes in the correlation structure and coactivity among multiple GABAergic subtypes, suggesting a broadly disorganized microcircuit functionality in mutant mice. Overall, we identify widespread and heterogeneous subtype-specific alterations in interneuron dynamics during spatial reward navigation, reflecting impaired flexibility and organization in CA1 inhibitory microcircuits. Our study provides critical insights into how schizophrenia-risk mutations affect local-circuit interactions among diverse cell types in the mouse hippocampus during learning and spatial navigation.

## INTRODUCTION

Adults and children with the 22q11.2 deletion syndrome (22q11.2DS) exhibit social and emotional impairments alongside a range of cognitive deficits^1,2^. The 22q11.2 deletion is one of the strongest genetic risk factors for schizophrenia, leading to a marked ∼30-fold increase in the risk of developing the disorder^1,3,4^. Cognitive deficits represent core features of schizophrenia and are likely mediated by the same abnormal physiological processes that underlie other more prototypical symptoms^5–8^. Schizophrenia patients experience a spectrum of cognitive deficits, which are often treatment-resistant. Among these, episodic memory dysfunction^5,9,10^ is both relatively common and highly debilitating. Episodic memory deficits, which show some of the largest effect sizes among cognitive impairments in schizophrenia^11^, are often present prior to the onset of psychosis, and are frequently observed in relatives of affected individuals^11–13^. Episodic memory relies on the hippocampus, which exhibits both anatomical and functional deficits in schizophrenia and 22q11.2 deletion patients^14–22^.

While mouse models cannot fully capture the symptoms associated with psychiatric diseases, they are valuable for modeling disease-associated mutations and assessing core aspects of a disorder that manifest in an equivalent manner in rodents^6,23,24^. The hippocampus is one of the most evolutionarily conserved brain regions^25,26^ and the mouse hippocampus recapitulates human features in both normal physiology^27–29^ and pathophysiology of neurological disorders^30–33^. By using animal models of rare, highly penetrant risk mutations, and coupling these models with behavioral tasks that assess key cognitive features of learning and memory, researchers can apply high-resolution neuroscience techniques to gain unprecedented insights into the mechanisms underlying mental illness^34–36^. These approaches have introduced a new biological framework for studying psychiatric disease, emphasizing disordered activity and plasticity across both local and long-range neuronal assemblies and circuits^34,35,37–47^. Such disruptions may reveal common points of convergence of genetic liability, leading to erratic neuronal activation patterns that trigger the constellation of cognitive symptoms, including episodic memory deficits^23^.

Hippocampal place cells, a subset of hippocampal principal cells that are selectively active in specific environmental locations, provide cellular substrates for episodic memory and goal-directed spatial learning^48,49^. Place cell ensembles are both sparse and dynamic^43,50–57^, forming rapidly in novel environments^58–60^, and undergoing experience-dependent stabilization as the environment transitions from novel to familiar^61–63^. Place cells are also responsive to salient reinforcers, such as rewards^43,51,64–67^, with distinct, sparse populations of place cells representing different contexts^52,63,67–69^. Our previous study uncovered behavioral and place cell correlates of episodic memory deficits in *Df(16)A^+/−^* mice, an animal model for the 22q11.2 deletion^70^, by using a spatial goal-oriented, hippocampal-dependent learning task^43^. *Df(16)A^+/−^* mice exhibited a marked impairment in learning performance when both the contextual cues and reward locations were changed during the experimental paradigm. Furthermore, place cells in the dorsal hippocampal area CA1 of mutant mice exhibited reduced long-term stability, impaired context-related remapping, and a lack of reward-related reorganization. Despite these population-level deficits, slice electrophysiological recordings of CA1 pyramidal neurons failed to identify cell-intrinsic abnormalities, instead pointing to alterations in inhibitory input^71^.

These observations provide a crucial entry point for investigating how schizophrenia risk mutations lead to altered hippocampal microcircuit dynamics during learning. However, the role of many hippocampal circuit components has remained uncharacterized, representing a major gap in our understanding of circuit mechanisms that drive disease-predisposing genetic lesions into clinical symptoms. In particular, GABAergic interneurons (IN) play a crucial role in shaping the activity of excitatory neuron populations in the brain by regulating the timing, amplitude, and synchronization of neuronal firing^72–74^. CA1 pyramidal cell activity and plasticity are under stringent inhibitory control by a diverse array of IN subtypes, each with distinct morphological, molecular, and functional properties^75–77^. Different IN subtypes target specific subcellular compartments of excitatory neurons and have specialized roles in regulating microcircuit function^76,78–83^. Dysfunction in inhibitory circuits has been implicated in numerous psychiatric disorders, particularly in schizophrenia pathophysiology^41,42,84–93^. Therefore, understanding the role of inhibitory neurons in microcircuit dysfunction is key to unraveling the complexity of neural circuits and their contribution to various brain functions and psychiatric conditions. Yet, the *in vivo* alterations in activity dynamics across most CA1 IN subtypes resulting from 22q11.2 deletions remain completely unknown. Prior studies have largely examined IN subtypes in isolation (‘one cell-type at a time’)^87,94–96^, a limitation that precludes a circuit-level understanding of inhibition. This piecemeal approach has hindered the identification of disease-associated inhibitory dysfunction, highlighting the need for holistic experimental and analytical strategies that can simultaneously assess functional interactions among multiple IN subtypes.

To overcome the limitations of the previous studies, we innovate both in terms of applying advanced imaging methods and designing new analysis approaches to yield novel insights about hippocampal circuit organization. Namely, we utilized a three-dimensional two-photon (2p) Ca^2+^ imaging strategy with multiplexed molecular characterization to capture a simultaneous functional readout of a large volumetric population of GABAergic IN subtypes in the dorsal hippocampal CA1 area^71,72^. We then introduce a new computational model to quantify the interplay between molecular readouts of INs and their pairwise functional correlation patterns. This analysis enables us to approximate circuit co-activation changes amongst INs from different genotypes to make many-cell-types-at-time inferences about circuit differences (as opposed to existing one-cell-at-a-time approaches). These techniques were applied on *Df(16)A^+/−^*mice and wild-type (WT) littermates to address key gaps in our understanding of the local circuit alterations associated with the 22q11.2 deletion and to dissect the relationships between cellular components that contribute to the pathophysiology of schizophrenia-related memory deficits.

## RESULTS

### Interneuron subtype number or distribution in dorsal CA1 is unaltered in *Df(16)A^+/-^*mice

To assess interneuron dynamics within the hippocampus during navigation in *Df(16)A^+/-^* mice, we employed an unbiased, large-scale, 3D method for *in vivo* 2p Ca^2+^ imaging of dorsal CA1 INs with random access acousto-optical deflector (3D-AOD) microscopy (Fig. 1)^97,98^. We performed 3D-AOD 2p Ca^2+^ imaging in *Vgat-Cre::Df(16)A^+/−^*mice and WT littermates injected with a Cre-dependent adeno-associated virus for GCaMP7f expression in dorsal hippocampal area CA1 INs during a Random Foraging behavior paradigm (see Methods, Fig. 1A,B)^97,98^. After taking a Z-stack of the dorsal CA1 field of view (Fig. 1C), we manually selected individual IN somata distributed across the 3D volume (Fig. 1D). This approach enables simultaneous, longitudinal monitoring of activity dynamics in large IN populations (up to ∼250 cells per mouse) across all CA1 sublayers to effectively capture the local GABAergic microcircuitry (Fig. 1E). For the purpose of this study, we refer to the local CA1 region as a microcircuit of locally connected excitatory pyramidal neurons and inhibitory GABAergic interneurons, and we collected and analyzed the GABAergic IN components of this microcircuitry. We imaged the same INs for two consecutive days while the head-fixed mice navigated a spatially enriched treadmill belt for randomly deposited water rewards. We performed *post hoc* immunohistochemical (IHC) identification of imaged INs and manually registered them to their *in vivo* cell somas based on their GCaMP7f signal (Fig. 1F, Suppl. Fig. 2A-C). We identified five major IN subtypes: axo-axonic cells (AACs), parvalbumin-expressing basket cells (PVBCs), somatostatin-expressing cells (SomCs), ivy/neurogliaform cells (IvC/NGFCs), and cholecystokinin (CCKCs) cells (Fig. 1G, see Methods, Table 2)^75,76,82,97^. Other IN populations, including vasoactive intestinal polypeptide-expressing and calretinin-expressing interneurons, were not examined in this study due to the limitations in the number of IHC channels and antibody combinations available for use within the same animal. Our analysis revealed that the subtype-specific distribution of imaged INs across CA1 sublayers was not noticeably altered in the mutants, and there were no significant differences in the number of IN subtypes recorded per mouse or their proportional distribution within each CA1 layer (WT n=6 mice, 1127 total cells, 187.83 ± 15.22 cells per mouse; Df(16)A^+/-^ n = 5,775 total cells, 155 ± 30.21 cells per mouse) (Suppl. Fig. 1A-C). Using the markers in our subtyping strategy (see Methods), we quantified the total number of cells expressing Parvalbumin (PV+), Somatostatin (SST+), Neuropeptide Y (NPY+), Satb1+, and Cholecystokinin (CCK+) in dorsal CA1 and assessed labeled cell soma density by layer (Suppl. Fig. 1D-I). Due to limitations in imaging depth and resolution, fewer INs were recovered in deeper layers of the hippocampus. Unidentified INs, which represent a highly heterogeneous VGAT population with variable functional properties, were excluded from subsequent analyses. Consistent with findings from other mouse models of the 22q11.2 deletion^99^, none of these quantifications revealed significant differences in the number of immuno-positive cells for each marker.

**Figure 1.**
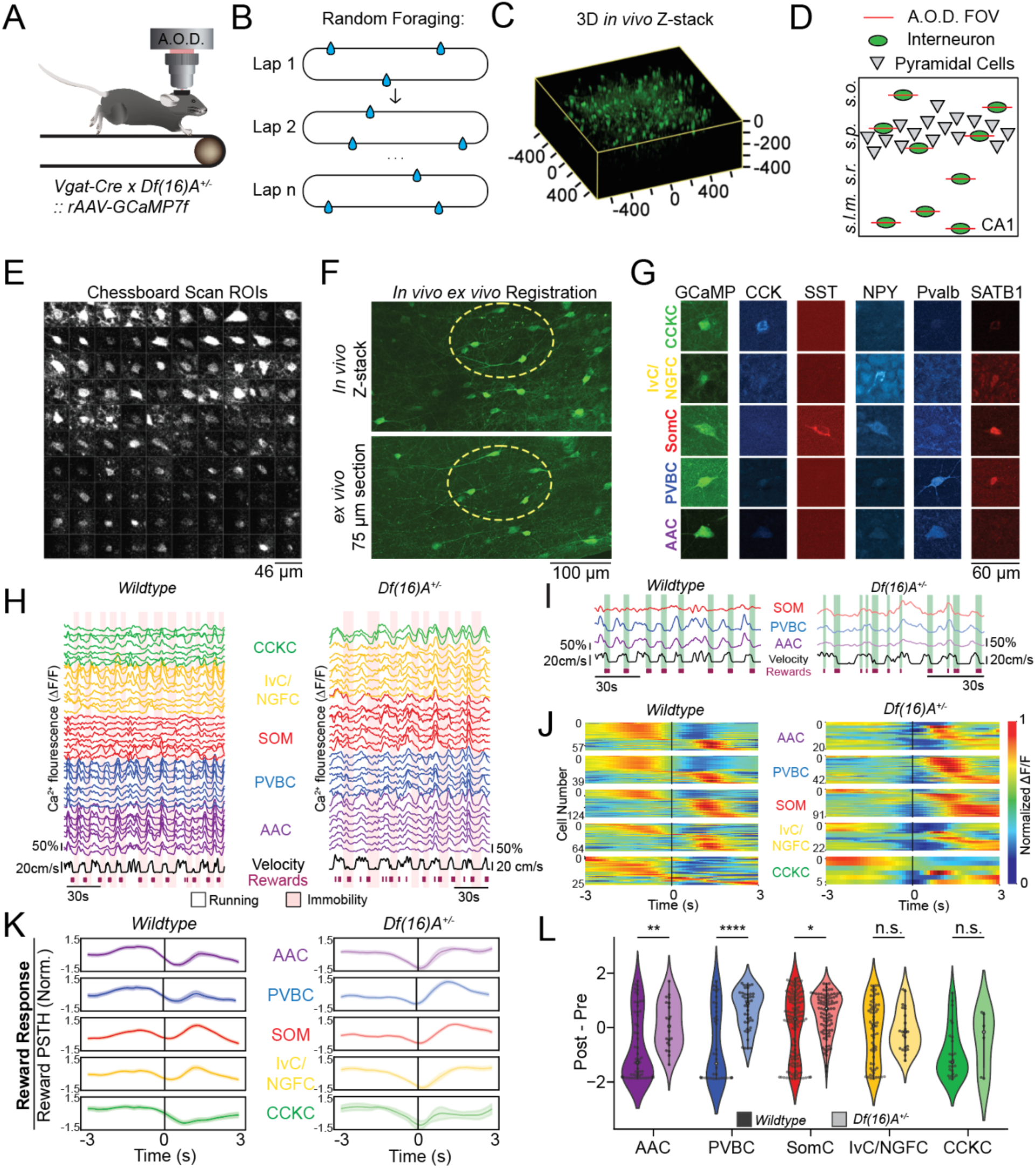
Df(16)A^+/-^ interneurons exhibit altered subtype-specific reward related responses. **A.** Schematic of the *in vivo* imaging setup using acousto-optical deflector (*AOD*) microscopy and viral GCaMP7f expression strategy. *Vgat-Cre::Df(16)A^+/−^* mice and WT littermates were injected with AAV1(*syn-FLEX-GCaMP7f*) in dorsal hippocampal area CA1 to express genetically encoded Ca^2+^ indicator, GCaMP7f, in interneurons (INs). **B.** Schematic of the Random Foraging behavior paradigm. Water-restricted mice were trained to run for randomly deposited water rewards across a 2-meter burlap belt with distinct features sewn across the belt. **C.** 3-dimensional reconstruction of *in vivo* z-stack of INs in CA1. **D.** Schematic of the 3D imaging targeting strategy on the AOD, coronal view. *s.o.: stratum oriens, s.p.: stratum pyramidale, s.r.: stratum radiatum, s.l.m.: stratum lacunosum moleculare*. **E.** Example time averages of the chessboard scan of 100 simultaneously imaged INs. *ROI*: regions of interest. **F.** Example pair of *in vivo* and *ex vivo* images manually registered between imaging modalities. **G.** Example images for each channel demonstrating the immunohistochemical (IHC) targeting strategy for IN cell-type characterization. Immunolabels are on the X-axis: *CCK*: cholecystokinin; *SST*: somatostatin; *NPY*: neuropeptide Y; *Pvalb*: parvalbumin. Cell-type labels used based on IHC strategy on the Y-axis: *AAC*: Axo-Axonic cells; *PVBC*: Parvalbumin Basket cell; *SomC*: Somatostatin-expressing cell; *IvC/NGFC*: Ivy/Neurogliaform cell; *CCKC*: Cholecystokinin-expressing cell. **H.** Example relative GCaMP7f fluorescence *(ΔF/F*) traces from molecularly identified interneurons during running and immobility bouts from 1 animal for each genotype. WT Cell Numbers: AAC = 62 cells, SomC = 132 cells, PVBC = 46 cells, CCKC = 32 cells, IvC/NGFC = 73 cells; *Df(16)A^+/-^* cell numbers: AAC = 34 cells, SomC = 125 cells, PVBC = 51 cells, CCKC = 14 cells, IvC/NGFC = 47 cells. **I.** Example relative GCaMP7f fluorescence *(ΔF/F*) select smoothed traces from interneurons highlighting rewarded areas during random foraging. **J.** Individual cell responses to reward during random foraging task. Left: WT; Right: *Df(16)A^+/-^*. **K.** Averaged z-scored reward-onset peri-event time averages (PETAs) of smoothed ΔF/F traces between −1.5 and 1.5 z-scores by cell type across all cells within cell type during the Random Foraging task for 3 seconds pre- and 3 seconds post-event. The shaded region represents the standard error. **L.** Post - Pre PETA responses per cell type before and after reward onset comparing the 3 seconds pre- to 3 seconds post-event. n.s.: not significant. *: P < 0.05. **: P < 0.01. ***: P < 0.001. See Suppl. Table 1 for detailed statistics.

### Unaltered locomotion-related interneurons dynamics in *Df(16)A^+/^*^-^ mice

To assess *in vivo* functional activity dynamics of molecularly identified CA1 interneurons (INs), *Vgat-Cre::Df(16)A^+/−^*mice and their wild-type (WT) littermates were water-restricted and trained to forage for randomly distributed water rewards on a spatially cued belt during 3D-AOD imaging (Fig. 1B). Individual movies were visually inspected from each preselected ROI per session based on the initial z-stack *in vivo,* motion corrected, and raw traces were extracted and processed for analysis (Methods, Suppl. Fig. 3). We smoothed the traces to focus on the slower components of calcium fluctuations in interneurons as there is limited ground truth data between calcium and spiking activity (Suppl. Fig. 3, 2^nd^ from bottom). Licking and average running velocity were consistent between *Df(16)A^+/-^*mice and WTs during this task (Suppl. Fig. 4A,B). Additionally, *Df(16)A^+/-^* mice ran the same number of laps without any changes in belt occupancy (Suppl. Fig. 4C,D). Most INs exhibited positive activity correlation with locomotion (Fig. 1H), which remained true in the described cell types in the mutant mice (Suppl. Fig. 4E)^97^. Similarly, CCKCs exhibited a bimodal correlation profile in both genotypes, with a subset of CCKCs showing negative velocity correlation^79,97^. Peri-event time average (PETA) analysis of subtype-specific IN activity during *Run-Start* and *Run-Stop* events showed no significant differences between genotypes (Suppl. Fig. 4F,G). Further analysis using a ridge regression model confirmed that velocity remained a significant predictor of IN activity, with no observable differences between the genotypes (Suppl. Fig. 4H). While licking was found to be a significant covariate for IN activity, the effect sizes were very small when compared with the disproportionately large impact of velocity. Thus, locomotion-related activity dynamics of INs remain largely unaltered in mutant mice.

### *Df(16)A^+/-^* interneurons exhibit subtype-specific alterations in reward-related inhibitory dynamics during random foraging

Locomotion equally drives interneuron activity in both genotypes (Fig. 1H, Suppl. Fig. 4). However, closer examination of activity traces suggested potential differences in IN responses to reward delivery between the genotypes (Fig. 1I). To isolate these differences, we smoothed the ΔF/F traces and regressed out the velocity component from the calcium traces to remove the highly correlated IN activity related to movement speed (Suppl. Fig. 3). We then investigated inhibitory responses to random rewards by analyzing neuronal activity during the random foraging task focusing on reward-onset PETAs. Our analysis revealed that more *Df(16)A^+/-^* INs exhibited stronger post-reward onset responses overall compared to wildtype INs (Fig. 1J). This post-reward enhanced activity was subtype-specific. For instance, SomCs, which typically showed increased post-reward activity in wildtype mice, exhibited an even greater recruitment in *Df(16)A^+/-^*mice. Similarly, AACs and PVBCs which typically exhibit decreased post-reward activity in WT mice, displayed a significant shift towards higher post-reward responses (Fig. 1K/L) in the *Df(16)A^+/-^* group. These findings indicate a substantial alteration in the inhibitory reward response to random rewards in Df(16)A+/- mice.

### *Df(16)A^+/-^* interneurons exhibit enhanced reward zone modulation and attenuated plasticity during a spatial navigation task

*Df(16)A+/-* mice exhibit hippocampal-dependent learning deficits^43,70,100–102^, consistent with what has been seen in human patients with 22q11.2DS^18,103–105^ and SCZ^106–109^. Considering our finding that inhibition is enhanced in response to random reward presentation and that *Df(16)A^+/-^* mice display decreased principal neuron enrichment near known reward zones^43^, we next aimed to characterize the spatial reward-related activity of INs, which is known to be cell type-specific^97,110,111^. To dissect how inhibition may play a role in poor cognitive flexibility during spatial navigation without the confound of a completely altered cognitive state (ie, not understanding the task or seeking vs finding the reward location), we designed a simple spatially guided reward translocation task. We trained the mice on a fixed reward task for 3 days prior to imaging with a non-operantly delivered water reward (Fig. 2A). On the fourth day, CA1 IN Ca^2+^ activity was imaged during a 10-min session in the familiar condition (Fig. 2B). On the next day, the location of the non-operant reward was translocated to another position on the treadmill belt and was repeated at that new position on the third day (Reward Translocation task, Fig. 2B). Both wildtype and *Df(16)A^+/-^* mice performed this task as expected, as evidenced by increased anticipatory licking leading up to and inside the reward zones (Suppl. Fig. 5A). Expectedly, the mice displayed a quick adjustment to the translocated reward zone based on their licking behavior on the first translocation day (Day 2) as the water is automatically given once they enter the reward zone (Suppl. Fig. 5A, middle). *Df(16)A^+/-^* mice exhibited more frequent licking overall than their WT counterparts (Suppl. Fig. 5A/B). When we examined the total licks per day, we found that this effect was largely driven by the first day, and that this significant difference is lost after the reward zone is translocated (Suppl. Fig. 5C). This phenotype was also specific to the fixed reward tasks, as there were no differences in licking behavior during the Random Foraging task (Suppl. Fig. 4A), indicating that hyperactivity in this mouse line^70,102,112^ was unlikely to explain these findings. When comparing all three days together, we additionally found that the mutant mice spent a greater proportion of time in the reward zone on the second day at the new reward zone and licked more within the reward zone (Suppl. Fig. 5D/J/K/L) despite running the same number of laps (Suppl. Fig. 5G) and having equivalent average speed (Suppl. Fig. 5H/I). However, when examining licking behavior within the reward zone and in anticipation of the reward zone (10 cm/5 bins prior), a measure of task learning, there was no significant difference between genotypes in either total licking or the proportion of anticipatory licking (Suppl. Fig. 5F,G). Overall, both WT and *Df(16)A^+/-^* mice navigated the spatial navigation task well while *Df(16)A^+/-^* mice exhibited subtle changes in behavior before and after the reward zone was translocated.

**Figure 2.**
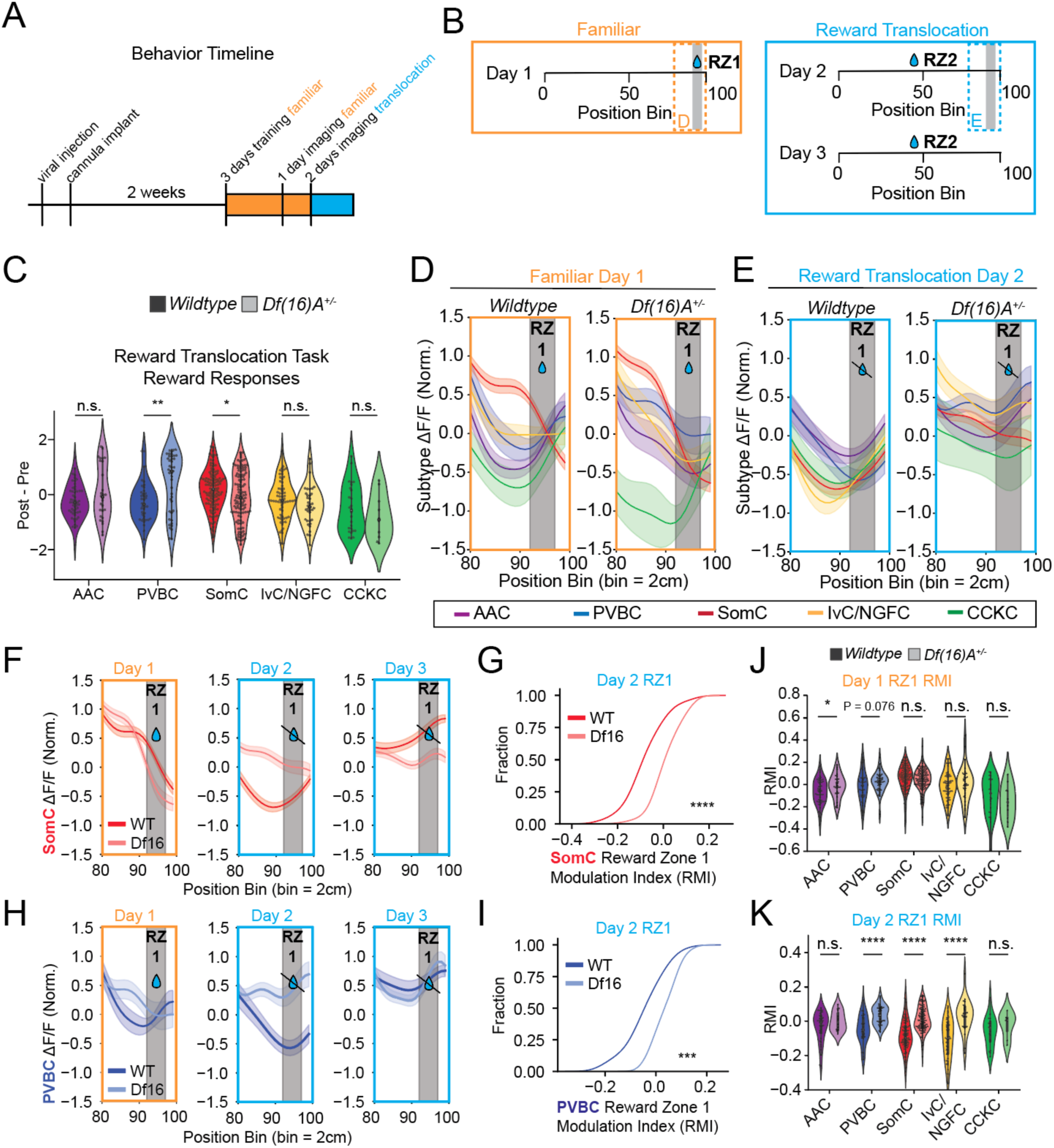
Aberrant subtype-specific activity during spatial reward navigation task. **A.** Reward translocation task timeline schematic. Orange: familiar task session days, Blue: reward translocation days. **B.** Reward translocation task (*RZ*: reward zone) schematic for the three days of imaging, the familiar reward zone days (top), and the two reward translocation days (bottom). **C.** Post - Pre cell responses centered around the reward onset for reward translocation task observed across all three days. **D.** Average responses of each cell type in the preceding 40 cm of the belt to the reward location in the WT (left) and *Df(16)A^+/-^* (right) on Day 1. **E.** Average responses of each cell type in the preceding 40 cm of the belt to the prior reward location in the WT (left) and *Df(16)A^+/-^* (right) on Day 2 after reward translocation. **F.** Average responses of SomCs in the preceding 40 cm of the belt to the old reward location in the WT (red) and *Df(16)A^+/^*^-^ (pink) on all three recording days. **G.** Reward zone modulation index (*RMI,* see Methods under Reward Modulation Index) for SomCs at the old reward zone after reward translocation. **H.** Average responses of PVBCs in the preceding 40 cm of the belt to the old reward location in the WT (dark blue) and *Df(16)A^+/-^*(light blue) on all three recording days. **I.** RMI for PVBC cells at the old reward zone after reward translocation. **J.** Reward modulation index in Reward Zone 1 on Day 1 per cell type. **K.** Reward modulation index in Reward Zone 1 on Day 2 per cell type. n.s.: not significant. *: P < 0.05. **: P < 0.01. ***: P < 0.001. See Suppl. Table 1 for detailed statistics.

We next interrogated distinct reward-related inhibitory responses during the reward translocation task (Fig. 2B) and observed subtype-specific changes in reward-related activity of multiple IN subtypes between genotypes. By comparing post-pre reward responses from reward-onset PETAs (Suppl. Fig. 4I) we identified increased pre-reward responses from SomCs and a stronger post-reward response from PVBCs (Suppl. Fig. 6A/B, Fig. 2C). We next examined the spatially defined average activity of each IN subtype preceding known reward locations. As expected, SomC activity was enriched leading up to both the familiar (Fig. 2D) and translocated reward zones in both the wildtypes and *Df(16)A^+/-^* INs (Suppl. Fig. 6C)^97^. However, in addition to SomCs, other IN subtypes, including PVBCs and AACs, also showed increased activity leading up to reward zones in *Df(16)A^+/-^*INs (Fig. 2D, Suppl. Fig. 6C, right). After reward translocation, we observed that SomCs and PVBCs maintained elevated activity levels preceding the old reward zone, indicating a persistent, inflexible response to prior reward locations in the mutant mice (Fig. 2E-H). To quantify these findings, we computed a reward zone modulation index (RMI, as previously described^97^) which normalized pre-reward zone activity to activity levels across the remainder of the belt (Fig. 2G/I, Suppl. Fig. 6D/E). This analysis confirmed sustained SomC, PVBC, and IvC/NGFC activity in the old reward zones of *Df(16)A^+/-^* mice compared to their WT littermates after reward translocation (Fig. 2G,I,K). At the new reward zone, while *Df(16)A^+/-^*SomCs enrich preceding the reward zone on the first day of the context switch (Suppl. 6C), they are significantly less enriched than the WTs (Suppl. 6F). This enrichment is flipped on the second day of the reward translocation task, with SomCs, PVBCs, AACs, IvC/NGFCs and CCKCs all being more enriched preceding the reward zone compared to the WTs. Interestingly, AACs uniquely exhibited aberrant reward-related activity only in the active reward zones (Fig. 2D, J, Suppl. Fig. 6C, F), whereas IvC/NGFCs showed aberrant activity only after the reward had been translocated (Fig. 2J/K). Of note, we did find that WT SomCs exhibit a significant RMI difference on Day 3 preceding the old reward zone compared to *Df(16)A^+/-^* SomCs; it is our interpretation that this is activity modulated by the new reward zone further down the belt, as this increased activity does not actually peak preceding the old reward zone, but after, unlike all other forms of reward-related enrichment described here ((Fig. 2D, 2E, 2F (left, middle), Fig. 2H (left, middle)). Overall, these findings identify task-dependent reward-related alterations in the *Df(16)A^+/-^* CA1 interneuron activity, reflecting both enhanced and inflexible local inhibition during spatial navigation.

### *Df(16)A^+/-^* interneurons represent reduced spatial information across tasks

We previously identified impaired place cell stability in CA1 and fewer place fields of place cells in mutant mice^43^. Given that place field formation and stability are under strong local inhibitory control^64,113,114^, and a subset of INs themselves exhibit positive or negative place fields in WT mice^83,97,98,113,115,116^, it is plausible that the aberrant reward-related activity exhibited by *Df(16)A^+/-^*INs (Fig. 1, 2) could fail to adequately support place cell spatial representations and contribute to spatial learning deficits previously observed in *Df(16)A^+/-^* mice^43^. To investigate this, we specifically examined spatial IN activity by asking how well we could decode the mouse position during the Random Foraging task based solely on IN activity (Fig. 3A). We trained a support vector machine based on all recorded IN activity (see Methods) and found a significant increase in decoding error in mutant mice INs compared to controls. This deficit persisted even after correcting for the number of cells recorded per animal and the number of laps run by mice (Fig. 3B,C). In addition, we also observed a non-significant decrease in spatial information encoded in mutant IN activity as measured by the Skaggs Information content (Fig. 3D)^117,118^.

**Figure 3:**
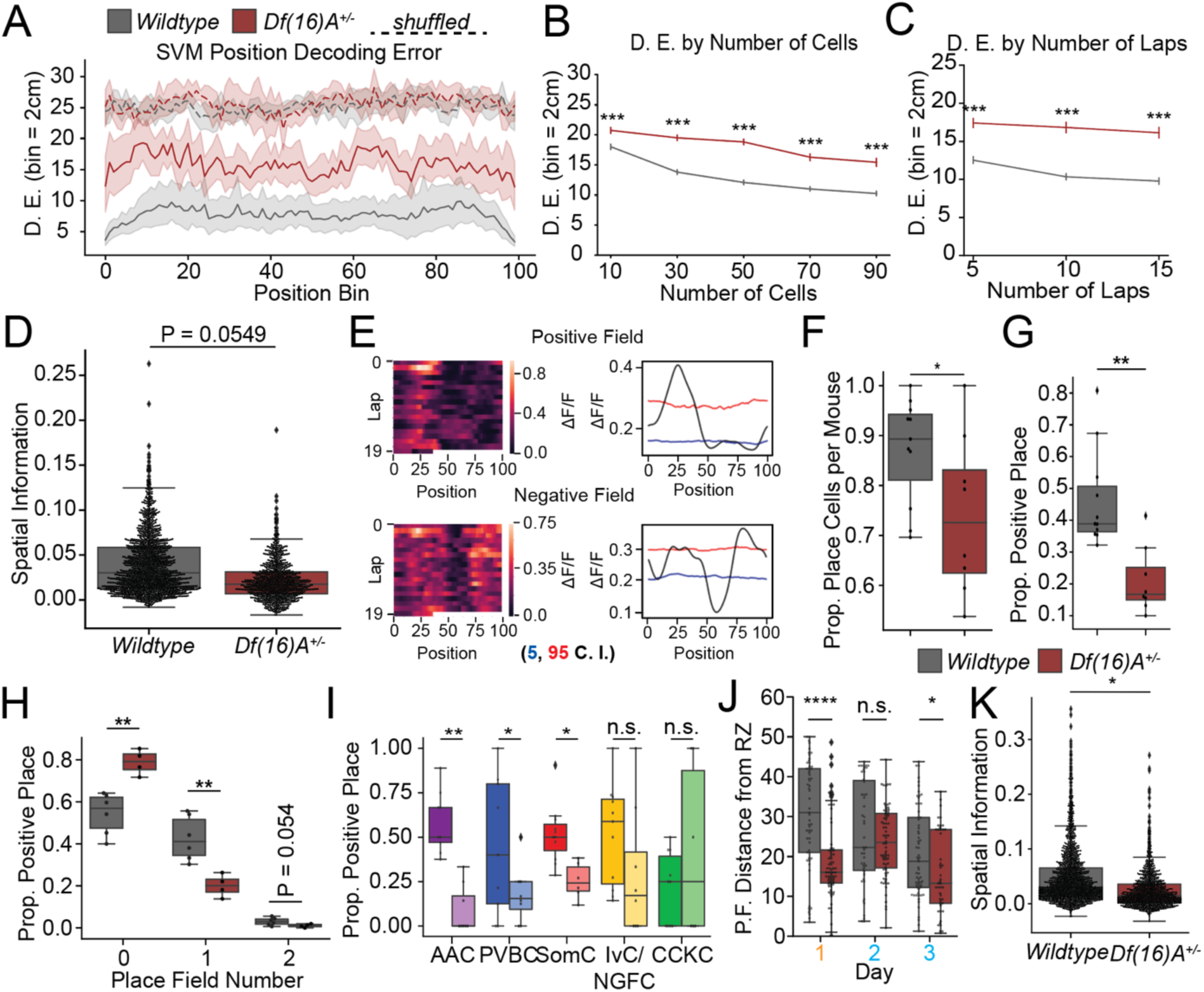
Reduced spatial information in *Df(16)A^+/-^* interneuron activity. **A.** All recorded interneuron (IN) activity was used to train a support vector machine (*SVM*) to decode discrete position bins (*D.E.*: decoding error) across the belt. Each bin is equivalent to 2 cm. **B.** SVM results iterated by the number of cells used with constant laps (trials) per genotype per bin. **C.** SVM results iterated by the number of laps (trials) run with a constant number of cells per genotype per bin. **D.** Skaggs information content of IN activity by genotype. Dots represent individual cell information content. **E.** Example lap-by-lap activity (left) and smoothed, averaged traces (right) and for cells with a positive place field (top) and a negative place field (bottom) based on the Peak Place method. Blue lines: lower bound to define significant negative place fields, red lines define significant C.I.: confidence interval. **F.** Place field proportion between genotypes. **G.** Proportion of positive place fields detected. **H.** Number of positive place fields detected per cell. **I.** Place field proportions by interneuron cell type. **J.** Distance from the center of the detected positive place field (P.F.), surpassing the 95th percentile shuffle distribution) to the center of the reward zone. **K.** Skaggs information content of interneuron activity by genotype during the reward translocation task. n.s.: not significant. *: P < 0.05. **: P < 0.01. ***: P < 0.001. See Suppl. Table 1 for detailed statistics.

Next, we calculated spatial tuning curves for spatial selectivity in all IN subtypes during random foraging sessions using the peak method (Fig. 3E)^97,119^, and found that IN place field widths did not differ between genotypes (Suppl. Fig. 7A). Most place fields were detected within 30 bins of the track (Suppl. Fig. 7A). However, we detected fewer INs with place fields in *Df(16)A^+/-^*INs compared with controls (Fig. 3F) and this deficit became more pronounced when cells with exclusively negative place fields were excluded (Fig. 3G, Suppl. Fig. 7B). Notably, we did not see a significant reduction in total negative place fields between genotypes (Suppl. Fig. 7C), suggesting a specific loss of positive place field information in mutant IN activity. When considering all detected place field types, mutant INs appear to generate fewer place fields per cell (Suppl. Fig. 7D). This deficit was particularly pronounced when again focusing solely on cells with positive place fields (Fig. 3H), indicating a selective loss in positive place field enrichment in the mutant INs. Further analysis of cell types involved in this reduction revealed that AACs, PVBCs, and SomCs exhibited significant reductions in the proportion of positive place cells, with IvC/NGFCs potentially showing decreased place preference as well l (Fig. 3I). Overall, fewer INs exhibit spatial selectivity in mutant mice (Fig. 3).

Finally, we assessed if the fixed reward zones in the Reward Translocation task resulted in increased spatial representation between the genotypes. When accounting for spatial representations of reward, there was no significant difference in the number of place fields detected using the peak method during this task (Suppl. Fig. 7E). However, when we examined the spatial relationship of these fields regarding reward by computing the distance between the center of each positive place field and the reward zone, we found that the positive IN place fields were significantly closer to the reward in mutant mice when compared to littermate controls (Fig. 3J) both before and after the reward is translocated. This trend is only lost on the first day of translocation (Day 2) while IN activity enrichment at the old location persists. Furthermore, consistent with our findings in the Random Foraging task when comparing all three days, there was still a significant decrease in spatial information represented in mutant IN activity during the Reward Translocation task (Fig. 3K) as measured using the Skaggs Information content method^118^. This suggests that despite the aberrant reward-related activity observed in PVBCs, AACs, and IvC/NGFCs, this altered activity did not contribute to an increase in the overall spatial information retained from the treadmill belt. Instead, it led to severely altered and persistent reward-related activity (Fig. 2).

### Subtype correlation structure is weakened in the *Df(16)A^+/-^* CA1

The distributed IN subtype alterations observed during spatial navigation for rewards suggest a potential reduction in coordinated inhibitory dynamics within the *Df(16)A^+/-^* hippocampal CA1 microcircuit. Microcircuit coordination is essential for appropriate information flow and network stability^113,114^. Leveraging our ability to image multiple IN subtypes simultaneously, we investigated whether the coordinated activity patterns in dorsal CA1 INs are altered during spatial navigation tasks (Fig. 4A). We compared activity between subtypes within each genotype using Pearson’s cross-correlation analysis of each cell within a type to the average activity of a given cell class during a given session for each task and genotype. For each matrix, comparisons were done with the cell type on the left to the cell types on the bottom, resulting in a “directional” analysis that is asymmetric due to normalization when aggregated across animals. This asymmetry is expected as we investigate how each cell type relate to others across animals with differing numbers of cells per subtype. In wildtype mice, IN subtypes exhibit highly correlated activity patterns (even after normalizing for mouse velocity) recapitulating known functional relationships between subtypes (Fig. 4A, 4B/D left). For example, the CCKC subtype is comprised of both perisomatic-targeting basket cells (CCKBCs) and dendrite-targeting CCK+ INs^79,82,120^. CCKBCs exhibit uniquely anticorrelated activity with velocity^79,97^, and as expected, we found that CCKCs in WT mice showed the lowest within-group and cross-group correlations of any IN type characterized in this study (Fig. 4B, left). At the CA1 microcircuit level, correlation structures differ significantly between genotypes across both tasks (Fig. 4C/E, Suppl. Table 3). While both genotypes exhibit altered correlation patterns between tasks as well (Fig. 4D, Suppl. Table 3), the directionality of the correlations diverged: WTs exhibit increased CA1 GABAergic microcircuit correlation structure in the RT task compared with the RF task, whereas the *Df(16)A^+/-^*microcircuit exhibited decorrelation. Significant reductions in the correlation structures across subtypes during the RT task portray a global loss of coordinated microcircuit activity rather than implicating a specific subtype (Fig. 4E), whereas SomC correlated structure alone is significantly altered during random foraging (Fig. 4C).

**Figure 4:**
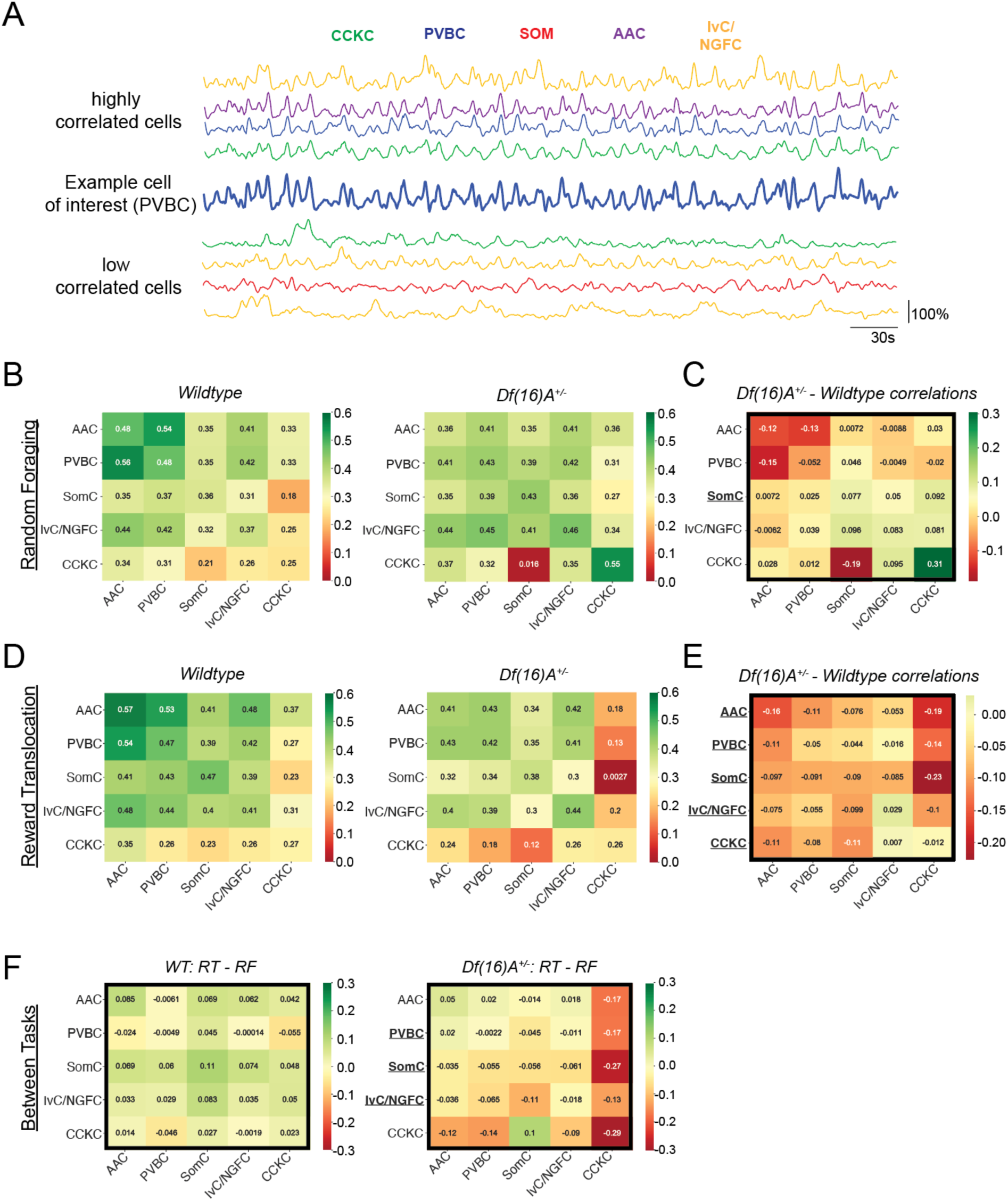
Coordinated activity is diminished in the *Df(16)A^+/-^* CA1 microcircuit. **A.** Example smoothed and velocity regressed traces of cells with visibly high correlation (top) and low correlation (bottom) with the PVBC trace in the middle. **B.** Pairwise Pearson’s correlation coefficient value matrix between cell types of (Δ*F*/*F*) traces across the session during the Random Foraging paradigm from WT (left), *Df(16)A^+/-^* (right) and the **C.** comparative differences between genotypes. **D.** Pairwise Pearson’s correlation coefficient value matrix between cell types of (Δ*F*/*F*) traces across the Reward Translocation sessions from WT (left), *Df(16)A^+/-^*(right) and the comparative differences between genotypes **E.**. **F.** Comparative differences within groups (genotypes) between tasks. Bolded outlines indicate significant microcircuit differences, bolded and underlined cell type names indicate significant cell type correlation pattern differences. *: P < 0.05. **: P < 0.01. ***: P < 0.001. See Suppl. Table 3 and Suppl. Fig. 8 for detailed statistics and visualization.

### A probabilistic graphical model uncovers altered coactivity patterns between interneuron subtypes in *Df(16)A^+/-^* mice

The loss of coordinated activity in *Df(16)A^+/-^* INs hints at alterations in cell-cell connectivity or responses to a common input (hereafter referred to as *coactivity*). We analyzed the coactivity patterns between IN subtypes across sessions, animals and behaviors to infer whether there are consistencies across biological replicates (Suppl. Fig. 9). Further, differences in consistency between different genotypes in the coactivity patterns between INs were used to suggest an increase or decrease in organization of the circuits that these INs were part of. For example, if the coactivity between SomC’s and all other INs was consistent and predictable across most of the WTs but no longer predictable in *Df(16)A^+/-^ mice*, we would deduce that circuit organization in *Df(16)A^+/-^* mice was not driven by these cell types, and therefore relatively disorganized. To quantify all of these effects and signatures of coactivity between INs, we devised a probabilistic graphical model (Suppl. Fig. 9A-D, see Methods) to model cell-types as a function of their co-activation patterns with all other IN subtypes in CA1 (Fig. 5A). The graphical representation of this model (Suppl. Fig. 9A) illustrates the hierarchical nature of the model, highlighting the dependencies between the different levels of interaction. For example, we model that each cell has a correlation value with all other cells in our observed dataset (Suppl. 9D). Then, we capture the coactivation of this cell with different cell-types as the maximum absolute correlation value amongst all cells of the specified type. This captures the most likely coactivation partner of a cell within each of the IN subtypes (thus, each cell is represented by a vector of K coactivations for K IN subtypes). We term this cell-to-cell-type coactivation (e.g., single-cell vs. all IN subtypes). We then take averages of these cell-to-cell-type values for all cells within a cell-type to deduce the expected *cell-type-to-cell-type* co-activations (e.g., SomC vs. all IN subtypes). Note that our cell-to-cell “co-activation” value is critically different than the average correlation between cell types that are used in the analysis of the previous section. We posit that the average of maximum absolute correlations (termed co-activation here) will capture interactions between cells than average correlations (used in the Fig. 4 analysis). All these interactions, both *cell-to-cell-type,* and *cell-type-to-cell-type* are modeled under a hierarchical Bayesian probabilistic model where we infer parameters of these distributions (e.g., mean and covariance of *cell-type-to-cell-type* co-activation patterns) to make quantitative and probabilistic statements about observed data. Using this model, we aimed to achieve two goals: 1) assess how co-activation profiles changed in mutant mice to infer microcircuit alterations and 2) determine the predictability of the molecular identity of INs based on their interaction profiles with known IN subtypes.

**Figure 5:**
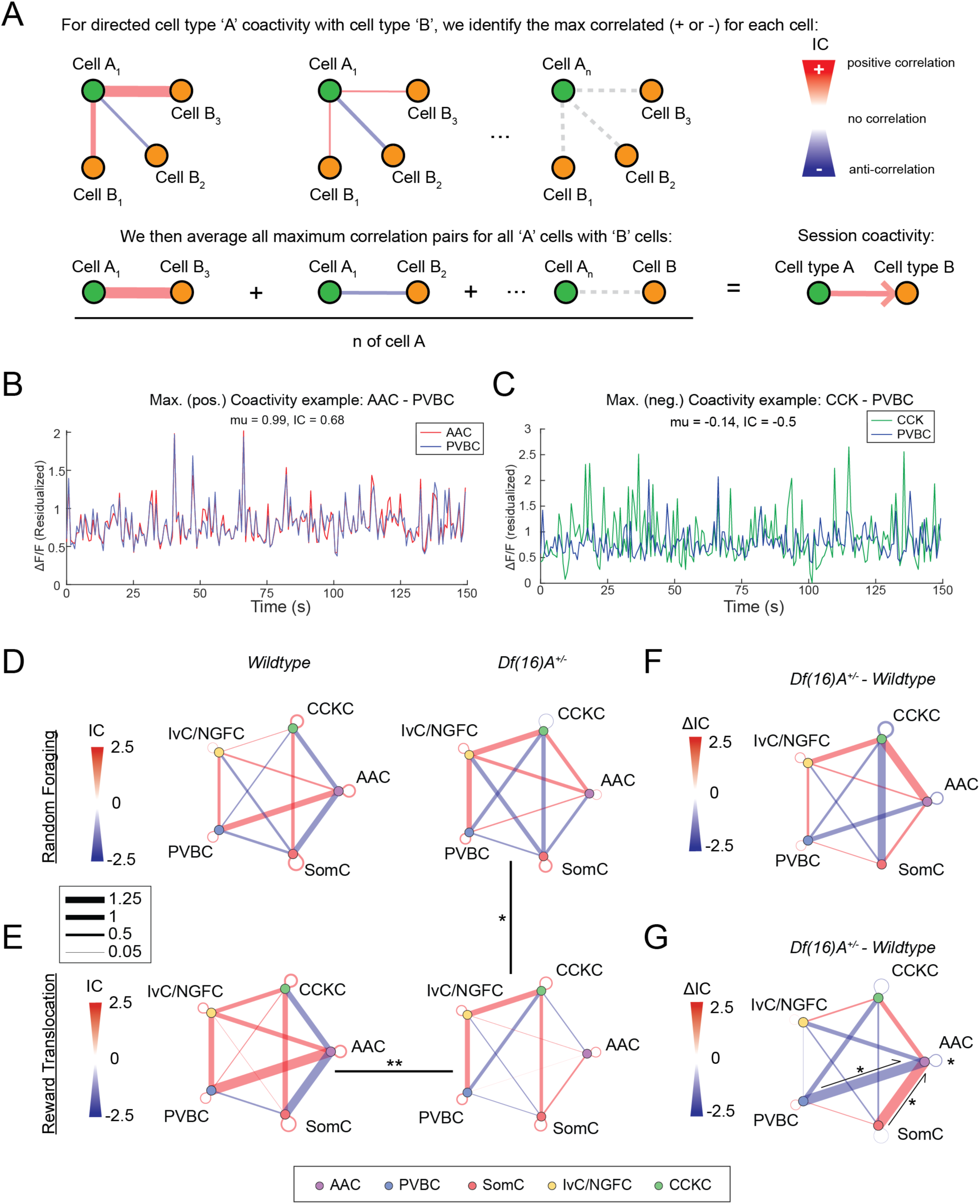
Coactivity alterations are genotype and task dependent. **A.** Schematic depicting how the model identifies the average maximum directed connection between a given pair of cells to estimate coactivity motifs. **B.** Example positive maximum coactive cells and **C.** example anticorrelated maximum coactive cells. (D/E) Circuit representation of the cell-to-cell-type interactions of Wildtype (left) and *Df(16)A^+/-^* (right) interneurons and by behavior paradigm (top: **D.** Random Foraging, bottom: **E.** Reward Translocation task), as determined by the probabilistic graphical model. Nodes represent different interneuron types, with colors corresponding to AAC (purple), CCKC (green), IvC/NGFC (blue), PVBC (yellow), and SomC (red). Edges connecting the nodes reflect the z-scores of the differences between the average correlations, with red edges indicating a correlation greater than expected by chance and blue edges indicating less than expected. **F.** and **G.** Circuit representation of the difference in cell-to-cell-type interactions between mutant *Df(16)A^+/-^* and WT genotypes by behavior paradigm (top: Random Foraging, bottom: Reward Translocation task), as determined by the probabilistic graphical model.

To probe cell-type activity relationships during behavior, we visualized the average cell-type interactions (*Ω*) to reveal the cell type-functional correlations within the hippocampal network. Similar to in Figure 4, comparisons are directional, but for simplified visualization, a single weighted line is used based on the arithmetic average between the two cell types. Directional arrows are included to illustrate significant effects between cell types, and all directional statistics are included in Supplementary Table 4. This notably recapitulated known mutual positively correlated (Fig. 5B) and anticorrelated interactions such as those between CCKC and PVBC type INs during Random Foraging (Fig. 5C, 5D left)^79,97,98^. In WT mice, most cell type relationships tended to be more coactive (or anti-coactive) than expected under the assumption that their activity was completely random during both behavioral tasks (Fig. 5D/E, left). When we compared the relative coactivity patterns between genotypes, we found that while there is no significant difference between genotype patterns during the Random Foraging task, there is a difference when comparing genotypes in the Reward Translocation task. This change seems to be driven more by changes in *Df(16)A^+/-^*coactivity, as *Df(16)A^+/-^* coactivity also differs between tasks (Fig. 5D/E, Suppl. Table 4). Notably, the change in coactivity between genotypes in the Reward Translocation task seems to be primarily attributed to changes during the first day of the reward translocation (Day 2) (Suppl. Fig. 10, middle, Suppl. Table 4). At the cell-to-cell comparison level, this effect may in part be driven by the sign flip of PVBC coactivity with AACs, which is typically a high positive coactivity pair in the WTs (Fig. 5B, 5D/E left, Suppl. Fig 10.) Overall, changes in the coactivity patterns seem to be driven by more global microcircuit effects rather than cell type specific ones, as few cell to cell specific coactivations appear to be significantly altered on their own (Suppl. Fig. 10).

### Altered cell type predictability infers altered circuit organization in *Df(16)A^+/-^*

Neuronal cell-type is thought to be a key determinant of circuit function^79,97,121^. We investigated whether neuronal identity could be inferred from the co-activity patterns (introduced in the previous section) of IN subtypes in local hippocampal CA1 circuits of WT and mutant mice. The separation of INs along principal component space (i.e., SomC and AAC along PC1, CCKC against all other INs along PC2) in the low dimensional projection of interaction parameters (Suppl. Fig. 11C) suggests that our model effectively captures features of co-activation that facilitate cell type classification. To further assess the nature of these co-activity signatures, we attempted to decode IN cell types based on their co-activity patterns with other cell types. If a cell type is predictable from its co-activity patterns, we can infer that their co-activity pattern is stable across sessions and animals. As stated before, we used the relative predictability of cell-types from their coactivation patterns as a surrogate of how organized a circuit was. Our reasoning is that an organized circuit has consistent coactivation patterns across biological replicates (e.g., different animals), and thus, we can make accurate predictions of cell types from their coactivation patterns. In contrast, if a circuit is disorganized, then there may be a low amount of consistency of co-activation patterns across different animals, and thus predictive models will suffer in out-of-sample generalizability. Our probabilistic model demonstrated robust classification for all five IN types tested (AACs, CCKCs, PVBCs, SomCs, IvC/NGFCs) in the WT with accuracy significantly above random chance during both Random Foraging (Suppl. Fig. 11E, left) and Reward Translocation tasks (Suppl. Fig. 11F, left). In contrast, our results revealed both task-dependent and cell type-specific alterations in IN predictability in *Df(16)A^+/-^*mutants. During Random Foraging, neither AACs nor CCKCs were predictable, although the latter could be attributed to lower numbers of CCKCs recovered in the mutants (Suppl. Fig. 11E, right). Meanwhile, during the Reward Translocation task, all cell types remained significantly predictable. The fact that cell types remain predictable in the mutants implies that the alterations in coactivation are consistent across animals rather than arising from random circuit disorganization.

In conclusion, probabilistic graphical modeling of IN coactivity reveals task-dependent, genotype-specific reorganization of coordinated inhibitory dynamics across multiple IN subtypes (Fig. 4), supporting a model in which schizophrenia-associated genetic risk disrupts hippocampal microcircuit organization at the ensemble level rather than through isolated subtype dysfunction.

## DISCUSSION

Our study provides the first comprehensive characterization of inhibitory local circuit dynamics in the hippocampus of mutant mice carrying a schizophrenia-predisposing genetic lesion. We discovered that mice carrying the 22q11.2 deletion, one of the strongest genetic risk factors for cognitive dysfunction and schizophrenia, exhibit widespread disruptions in hippocampal IN dynamics and coactivation patterns during similar cognitive processing stages. By examining IN subtype-specific responses during distinct spatial navigation tasks, our study reveals distributed microcircuit-level alterations (Table 1). Importantly, these alterations are most fully captured when considering coordinated IN activity, and would not be fully revealed by analyses focused solely on individual IN subtypes.

**Table 1.**
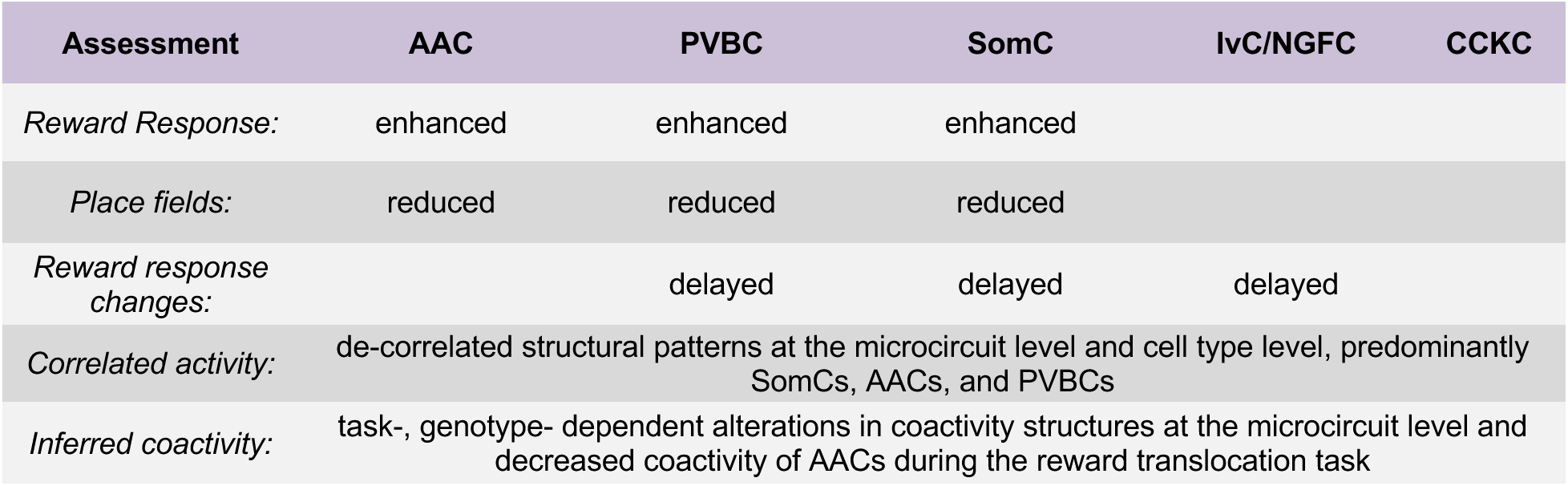
Summary of interneuron subtype-specific and microcircuit level deficits.

**Table 2.** Antibodies used in this study.

| Cell Marker | Round | Primary Antibody (dilution) | Catalog # | Secondary Antibody (dilution) | Catalog # |
| --- | --- | --- | --- | --- | --- |
| Cholecystikinin (CCK) | 1 | rabbit anti-proCCK (1:500) | Frontier Institute CCK-pro-Rb-Af350 | donkey anti-rabbit DyLight 405 (1:300) | Jackson 711-475-152 |
| Somatostatin (SST) | 1 | rat anti-somatostatin (1:500) | Millipore MAB354 | donkey anti-rat Alexa 568 (1:300) | Abcam ab175475 |
| Neuropeptide-Y (NPY) | 1 | sheep anti-NPY (1:500) | Abcam ab6173 | donkey anti-sheep F(ab)2 Alexa 647 (1:300) | Jackson 713-606-147 |
| Parvalbumin (Pvalb) | 2 | chicken anti-PV (1:5000) | Gift from Susan Brenner-Morton | donkey anti-chicken DyLight 405 (1:300) | Jackson 703-475-155 |
| SATB1 | 2 | rabbit anti-SATB1 (1:1000) | Abcam ab70004 | donkey anti-rabbit Alexa 647 (1:300) | Jackson 711-605-152 |

By monitoring IN activity dynamics during random foraging, we confirmed an overall positive velocity modulation of IN activity during locomotion in *both* WT and *Df(16)A^+/-^* mice. Despite the relative preservation of these fundamental features of hippocampal IN function^78,79,81,97,122–124^, our findings reveal pronounced alterations in the reward encoding across broad classes of INs in the *Df(16)A^+/^*^-^ mice compared to the WT littermates. Prior studies have implicated the role of specific IN subtypes, including SomCs and PVBCs^64,97,98,110,111^, in reward modulation. Consistently, we identified PVBCs and SomCs as exhibiting particularly altered reward modulation during both random foraging and fixed reward spatial navigation tasks, indicating pervasive disruption of the hippocampal reward-processing circuit. Previously, we showed that *Df(16)A^+/−^*CA1 pyramidal neurons display reduced enrichment in the reward zone during a goal-oriented learning task and mutant mice exhibit impaired extinction of licking at the previously rewarded location compared with WT littermates^43^. Taken together, these findings suggest that aberrant and inflexible subtype-specific IN activity may contribute to impaired reward-related representational dynamics and memory deficits in *Df(16)A^+/-^* mice.

Further, our results also uncovered a population-level reduction in the spatial information encoded by INs in the *Df(16)A^+/^*^-^ mice relative to WT littermates. Specifically, IN activity in the *Df(16)A^+/-^*mice carries significantly less spatial information, implying a weakened spatially localized inhibitory influence over CA1 pyramidal cell ensembles^97,114,125–129^. While positive place field detection in INs was reduced, negative place fields were preserved, indicating distinct and potentially dissociable roles for dynamic IN activity patterns^98^.

We identified large-scale relational networks of INs in *both* WT and *Df(16)A^+/-^* mice, revealing complex, correlational relationships between IN subtypes with selectively reduced correlations in the mutants. Through developing a probabilistic graphical model for coactivity, we extended these observations and uncovered significant, genotype-depended alterations in the coactivity patterns among IN subtypes, consistent with a less coordinated inhibitory microcircuit in *Df(16)A^+/-^* mice. Notably, our model effectively predicted several IN subtypes based solely on coactivity patterns, highlighting a distinct and discernible interaction pattern among IN classes in the mouse hippocampus. This modeling framework provides a foundation for future studies aimed at linking functional coactivity to underlying structural connectivity, potentially leading to a more comprehensive mapping of neural circuitry.

In the present study, we focused on major IN subtypes that directly target pyramidal cells in CA1. It remains to be determined whether the observed alterations in inhibitory subtype dynamics are primary drivers of the disrupted pyramidal cell activity we previously reported^43^ or, alternatively, whether they arise secondarily from altered recurrent or long-range excitatory inputs to INs^130–132^. Likewise, it will also be important to determine if disinhibitory circuits that target other INs^64,97,133–135^, including the subtypes investigated in the present study, also exhibit altered dynamics and contribute causally to the observed microcircuit dysfunction.

Studies in mouse models for schizophrenia and human patients have predominantly focused on PV-expressing INs, implicating reduced prevalence and activity of these cells in disorganized network activity^41,88,90,91,99,100,112,136–138^. Along these lines, destabilization in hippocampal neuronal dynamics analogous to the one described in Zaremba et al., 2017^43^ has been subsequently recapitulated in hippocampal *in vitro* preparations from another 22q11.2 deletion model (*Lgdel/+* mice) and correlated with deficits in the synchronized CA1 neuronal network activity, including diminished spike correlation between neuronal pairs and reduced theta frequency oscillations^112^. Importantly, these network and hippocampal-dependent behavioral deficits were partially or fully restored by acute pharmacological or chemogenetic manipulations of PV+ IN excitability, underscoring the role of inhibitory regulation in maintaining local circuit stability in the hippocampus. However, recent studies also implicate dysfunction in other IN types, including SomCs, Neuropeptide-Y-expressing INs, and AACs in various disease-related cortical circuits^7,45,89,135,139–145^. Importantly, these subtypes directly interact with one another, influencing microcircuit functional output. In agreement with this body of work, our results also show that PV+ INs dysfunction alone is insufficient to account for the full spectrum of inhibitory abnormalities associated with this high-risk schizophrenia locus. Diverse GABAergic INs express genes encompassed within the *Df(16)A^+/-^* deletion (Supp. Fig. 12A), with enrichment patterns that differ from those observed in glutamatergic neurons (Supp. Fig. 12B/C). Based on expression profiles, a network-wide molecular perturbation in INs could give rise to widespread dysfunction and altered excitation-inhibition balance rather isolated subtype-specific effects.

While we identified a limited behavioral phenotype in the reward translocation task, our work supports disrupted hippocampal GABAergic microcircuit dynamics as a candidate endophenotype associated with the 22q11.2 deletion syndrome. In human studies, endophenotypes linked to the 22q11.2 deletion syndrome have implicated neuronal dysfunction at both regional^146^ and whole-brain scales^147,148^. Given the complexity and heterogeneity of neurodevelopmental and psychiatric disorders, it is unlikely that a single deficit is sufficient to generate overt cognitive phenotypes^149–153^. Accordingly, identifying latent circuit-level dysfunction provides critical mechanistic insight into disease vulnerability. Our work highlights perturbations within a hippocampal structure central to cognitive function and motivates further investigation of inhibitory–excitatory microcircuits across brain regions. Moreover, given the variable penetrance and expressivity characteristic of psychiatric disorders, endophenotypes may serve as shared biological markers for therapeutic discovery and intervention^154–156^. While our study is observational, it establishes a technical and conceptual framework that enables future mechanistic interrogation using more demanding behavioral paradigms.

Overall, while further research is needed to fully elucidate the precise mechanisms by which 22q11.2 deletions and other risk mutations contribute to cognitive dysfunction and schizophrenia, it is increasingly clear that these effects are distributed across multiple IN subtypes within the brain (Table 1). This distribution affects both individual INs and ensembles of interacting IN cell types during cognitive operations in patients. Our findings suggest that genetic manipulation, modeling key aspects of schizophrenia risk such as 22q11.2 deletions, induces changes in the coactivity of local IN ensembles that are markedly more pronounced than alterations in individual INs. This implies that even seemingly small functional changes in individual INs can culminate in significant systems-level alterations when integrated into functional microcircuits.

Our study provides critical insights into how schizophrenia-predisposing mutations affecting cognitive dysfunction disrupt complex interactions among IN cell types in the mouse hippocampus during spatial navigation. It highlights a sophisticated interplay between disease-predisposing genetic lesions, the brain’s dynamic functional patterns, and, eventually, the manifestation of cognitive symptoms. Moreover, our findings have broader implications for the predictive modeling of neuron types based on their functional interactions. This approach offers a novel framework for understanding how the brain’s molecular structure and dynamic activity patterns are altered under pathological conditions. Future investigations will aid in expanding the molecular and structural correlates of the observed altered IN coactivity patterns, potentially leading to a more comprehensive understanding of how genetic liability leads to distorted maps of neural circuitry.

## METHODS

### RESOURCE AVAILABILITY

#### Lead Contact

Further information and requests for resources and reagents should be directed to and will be fulfilled by the lead contact, Attila Losonczy.

#### Materials Availability

The study did not generate new unique reagents or mouse lines.

#### Data and Code Availability

Data and code reported in this paper are available from the lead contact upon request.

### EXPERIMENTAL MODEL AND SUBJECT DETAILS

All experiments were conducted in accordance with NIH guidelines and with the approval of the Columbia University Institutional Animal Care and Use Committee. Experiments were performed with healthy, 2-5 month old, heterozygous adult male and female *Vgat-IRES-Cre* mice (Jackson Laboratory, Stock No: 016962, referred to as Vgat-Cre mice) that were crossed with hemizygous *Df(16)A^+/-^* mice generated as previously described^70^. Mice were held in a 12-hour light/dark cycle vivarium and housed 2-5 mice in each cage. All experiments were conducted during the light portion of the cycle. Most mice used in the experiment were male (n = 9), with (n =5 WT, n = 4 *Df(16)A^+/-^*) with n = 2 females (n = 1 WT, n = 1 *Df(16)A^+/-^)*. All wild-type mice were littermate controls of the *Df(16)A^+/-^* mice.

### METHOD DETAILS

#### Viruses

Cre-dependent recombinant adeno-associated virus (rAAV) for GCaMP7f (rAAV1-syn-FLEX-jGCaMP7f-WPRE, Addgene #1944820-AAV1, titer: ≥1:10^13^ vg/mL) were used to express GCaMP7f in the hippocampus of *Vgat-Cre* mice. The Cre-dependent recombinase enabled expression of Ca^2+^ sensor GCaMP7f in Vgat+ interneurons (IN)).

#### Viral injections and hippocampal window/headpost implant

During isoflurane anesthesia, *Df(16)A^+/-^* Vgat-Cre mice were placed onto a stereotaxic surgery instrument. Meloxicam and bupivacaine were administered subcutaneously to reduce discomfort. The skin was cut along the midline to expose the skull, and a craniotomy was performed over the right hemisphere with a dental drill. A Nanoject injector was loaded with a sterile glass capillary filled with mineral oil and rAAV1-syn-FLEX-jGCaMP7f-WPRE and lowered into dorsal CA1 of the right hemisphere. Injections were made at (in mm): AP −2.2, ML −1.75, DV - 1.8, −1.6, −1.4, −1.2, - 1.1, −1.0, −0.8 relative to bregma with 64 nl of virus injected at each DV level. The pipette was held within the brain for 5-10 minutes after the last injection and slowly raised from the brain. The skin was sutured, and a topical antibiotic was applied to the suture site. Four days after viral injection, a circular cannula with a glass window was inserted into the cavity produced by craniotomy as previously described in ref. ^130^. Similarly, as described above, mice were anesthetized with isoflurane and placed on a stereotaxic surgery instrument. Meloxicam and bupivacaine were subcutaneously given to the mice. The skull was exposed, and a 3 mm craniotomy was performed over the right hippocampus at AP −2.2, ML −1.75 relative to bregma, centered around the injection site. After removal of the dura, the cortex was slowly aspirated with negative pressure and the consistent application of ice-cold cortex buffer. When the horizontal fibers over CA1 were exposed, aspiration was stopped, and a sterile collagen membrane was applied to stop bleeding. A 3 mm circular cannula with a glass window was placed into the craniotomy and glued into the site with Vetbond. A headpost was then attached to the posterior of the skull with layers of dental cement. 1.0 mL of saline was subcutaneously given to each mouse, while the mice recovered in their home cage with a heat pad. All mice were monitored for 3 days of post-operative care until behavior training began.

#### Behavioral training and paradigms

Following the post-op recovery period, mice were acclimated to handling and head-fixation for a few days, while being water restricted to 85-90% of their original weight. Mice were trained to run on a two meter fabric cue-free belt for random and non-operantly dispensed water rewards that were decreased in number per lap as the mice increased the number of laps they ran per training session. The water reward amount per reward was also reduced in amount as lap number increased to encourage running. The water reward amount was calibrated per training session such that mice would gain 0.5 – 1.0 grams of weight at the end of a 10 minute behavior session running ∼15-20 laps. The lickport would be ‘open’ for 2 seconds once the mice entered the reward zone and they could receive water rewards operantly after the first non-operant reward for 2 seconds. Mice were trained for ∼1 week on the cue-free belt and then were transitioned to the fixed reward task on a spatially enriched belt.

#### Random Foraging Task

Mice were imaged on a two-meter spatially-enriched familiar context belt for 1 day with 3 random non-operant water rewards distributed across the length of the belt, randomized every lap. On day 2 of imaging, a novel belt was introduced with the same random reward paradigm. The belt consisted of four fabric ribbons with different textures and a variety of tactile cues: colored tape, foam strips, cotton pom poms, etc. The novel belt had distinct fabric materials and distinct tactile cues.

#### Reward Translocation Task

Mice were trained on a two meter cue-rich belt with a fixed non-operant water reward zone at the end of the belt for 3 or 4 days leading up to imaging. The belt consisted of four fabric ribbons with different textures and a variety of tactile cues: colored tape, foam strips, cotton pom poms, etc. The following day the mice were imaged with the familiar non-operant water reward zone location. On the second and third day of imaging, the reward zone was relocated to a new reward zone location at the center of the belt. The reward size was larger in this task than in random foraging task to account for the decreased number of rewards per lap.

#### Two-photon Ca^2+^ imaging

After the mice were acclimated and trained in the respective tasks, a high-resolution structural Z-stack was captured 24 hours prior to imaging, while the mouse was anesthetized with isoflurane. The reference Z-stack was used to identify the 3D position of the GCaMP-positive neurons for access by the custom-modified AOD (acousto-optic deflector) microscope (Femto3D-ATLAS). The AOD microscope was used as previously described ^97,98,113^.

A water immersion objective (x16 Nikon CF175) was used to focus at the dorsal CA1 pyramidal layer and then fixed in position. A tunable two-photon laser (Coherent Ultra II) was adjusted to wavelength = 920 nm. The high-resolution structural Z-stack was captured in CA1 from ∼100um below to glass to ∼600 μm to span the entirety of CA1 from the *stratum oriens* to the *stratum lacunosum-moleculare* with an 800×800 pixel capture with a resolution of 1.25 μm per pixel with a 4 μm step size. Laser power and photomultiplier tube (PMT) detectors (GaAsP, H10770PA-40, Hamamatsu) were compensated in power and gain throughout the Z-stack capture. After the Z-stack was completed, the mouse was returned to its home cage and given 24 hours to recover before functional imaging.

Using the integrated software from Femtonics (MES), the Z-stack was examined, and 100-150 INs were manually selected to generate a spreadsheet of the x-y-z positions at the center of each respective cell. These x-y-z positions served as the regions of interest (ROIs) that were then used on subsequent days by the AOD microscope to find the position of the corresponding interneurons. For each day of functional imaging in each mouse, the identical field of view was identified within the CA1 pyramidal layer, and then the ROIs were loaded using the software for AOD 3D imaging. The laser power is additionally compensated for z-depth across ROI coordinates during imaging as dictated by the parameters used during the initial z-stack to match the signal level between ROIs. Each imaging session was taken for 10 minutes, with 5-10 Hz for each recording experiment (frame rate depended on the number of ROIs, ROI size, imaging resolution, and scanning speed).

#### Immunohistochemistry

Mice were perfused with 20mL of Phosphate Buffered-Saline followed by 20 mL 4% paraformaldehyde (PFA, Electron Microscopy Sciences). The brains were stored in 4% PFA overnight and washed 3×5 min with PBS the next day. The brains were then sectioned for 75 μm transverse sections of the hippocampus on a vibratome and rinsed 3×15 min with PBS prior to immunohistochemical staining procedure. Histological tissue prep was performed as previously described^97,98^. In brief: sections were permeabilized 2×20 min with 0.3% Triton X-100 (Sigma-Aldrick), and subsequently blocked with 10% Normal Donkey Serum (Jackson ImmunoResearch) in PBST (PBS with 0.3% Triton X-100). After 45 minutes, the sections were incubated in PBS (containing 3 first-round primary antibodies, see below) for one hour at room temperature and then 2 days at 4°C with agitation. The sections were rinsed 3×15 min with PBS to wash out excess primary antibodies, and then, for two hours at room temperature, incubated in a PBS solution of secondary antibodies tagged for fluorescence (Table 2). Sections were then rinsed 5×15 min in PBS at room temperature and mounted onto rectangular glass slides with ThermoFisher Scientific aqueous mounting medium. After 1 hour of drying at 4°C, the slides were imaged. After imaging, PBS was used to remove coverslips, and sections were rinsed 3×15 min in PBS for round two of staining. The sections were blocked with 10% Normal Donkey Serum in PBST for 45 minutes before the addition of the round two primary antibodies. The secondary staining protocol (now using round two secondary antibodies) was then followed as completed in the first round. Staining for CCK, SOM and NPY in the first round led to stronger expression of fluorescence for those markers, opposed to staining for any of the three markers second (Table 2). Cells or sections with poor, unclear or incomplete staining were excluded from the analysis. For failed IHC staining for a particular marker in a section, all analysis for cell types requiring that marker were excluded from subsequent analysis.

#### Antibody information and staining strategy

##### Assignment of subtype identity based on immunostaining

Interneuron subtypes were assigned based on our previously published classification strategy^97^.

*Axo-axonic cells (AACs):*

Immunopositive for Pvalb and immunonegative for SATB1, SST, NPY, & CCK.

*Parvalbumin Basket Cells (PVBCs):*

Immunopositive for Pvalb and SATB1, immunonegative for SST, NPY, & CCK.

*Somatostatin-expressing cells (SomCs):*

Immunopositive for SST, with or without Neuropeptide Y (NPY) and SATB1, and immunonegative for Pvalb and CCK.

*Ivy / Neurogliaform cells (Ivy/NGFCs):*

Immunopositive for Neuropeptide Y, with or without SATB1, and immunonegative for SST, Pvalb, and CCK.

*Cholecystokinin-expressing cells (CCKCs):*

Immunopositive for proCCK, immunonegative for SST, PV, NPY and SATB1.

#### Confocal imaging

The NikonA1 confocal microscope used to take multi-channel fluorescence images (405 nm, 488 nm, 561 nm, and 640 nm excitation for the four far-red, blue, green and red colored channels) of labeled tissue sections. Image capturing yielded 2048 x 2048 pixel z-stacks of the entire depth of the section (with 3 microns between each image of a given stack). Z-stacks were viewable via Fiji (Image J).

#### Registration of confocal images to *in vivo* z-stacks

Unique groupings of cells from *in vivo* z-stacks were used as visual fiducial markers to manually register *ex vivo* 75 μm z-stack images in ImageJ, section by section. Care was taken to note the orientation of the *ex vivo* sections regarding the *in vivo* z-stack images. For registering the two imaging modalities to one another (*in vivo* and *ex vivo*), we had several advantages: The first being that the sparsity and distribution of interneurons using the VGAT-Cre + flox-depended GCaMP expression made it possible to easily see unique patterns of cells in all layers of CA1. Secondly, we took thick (75 uM) horizontal sections of the brain, almost perfectly in line with the plane of imaging *in vivo*, which facilitated relatively straightforward rough alignment (correct image transformation, ie flipping & turning) and fine alignment of cells (Suppl. 2A/B). Thirdly, the coordinates that we had generated from the initial *in vivo* Z-stack to image our interneurons chronically day after day were utilized in post processing for registration by mapping the imaged cell coordinates next to the cell number onto the Z-stack (depicted in Suppl. 2A-C). It is true that despite these advantages, slight angles in the tissue compared to the imaging plane *in vivo* would result in some cells being in or out of the same imaged planes. Because of our mapped coordinate references however, simply scrolling through the Z depth of both the immuno section or the *in vivo* Z-stack would recover these cells near perfectly (Suppl. Fig 2C). Thus, with this approach, we very rarely ever lost a cell due to registration drop out unless an *ex vivo* section was lost during processing.

##### Immunohistochemical marker density quantification with CATT

A novel Cell Analysis/Typing Tool (CATT) designed by E.V., was used to identify positive cells in confocal fluorescence microscopy images. The code uses the entry of pixel dimensions to generate an image that combines the four fluorescence color channels of an original section. A region of interest (ROI) was drawn to delineate distinct CA1 layers for the program to focus on counting cells within a certain hippocampal layer. Given the ROI and pixel dimensions, the program outputs the volume, as well as an image displaying all the IHC markers across the channels relative to the green channel (GCaMP7+ signal). Each positive cell marker was manually counted and divided by the ROI volume to derive positive marker density for a given ROI.

#### Behavior analysis

##### Licking behavior

Licks were detected via a capacitance sensor on the lickport depositing water rewards from our behavior controller system built in-house^157^ in the Losonczy lab. For each animal during each experiment, the two meter belt was divided into 100 bins (∼2cm per bin). The number of licks per bin per experiment were visualized to create licking curves. The proportion of licks that occur within the reward zone was also computed.

##### Reward zone occupancy

Each animal’s position per experiment was also digitized into 100 equal spatial bins representing the 2-meter belt. The occupancy time within the reward zone was quantified and compared between the genotypes.

#### Imaging analysis

##### Pre-processing of Ca^2+^ imaging data

Raw movies of each individual cell were independently motion-corrected using the whole-frame cross-registration implemented in the SIMA package^158^. We visually inspected each individual movie generated from each ROI: cells that exhibited particularly bright nuclei or nuclei that were brighter than the soma in GFP fluorescent signal during the imaging experiments were excluded from the analysis. Any cell that did not obviously exhibit dynamics was also excluded from the analysis. Signal did not differ across Z-depths (Suppl. Fig. 2D). ROIs were then hand-drawn over each imaged cell. Fluorescence was extracted from each drawn ROI using SIMA. ΔF/F was then computed based on a 1st percentile baseline value over a 30-second rolling window. The ΔF/F traces were then smoothed using a Savitzky-Golay filter with a window length of 9 and a polyorder of 3. For analysis involving velocity-regressed traces (Fig. 1, 2, 3), each animal’s velocity was smoothed with a Hanning filter with a window of 10, and then covariate linear regression was applied to each fluorescence trace where the smoothed velocity was regressed out (Suppl. Fig. 3).

##### Velocity correlation

For each cell, velocity correlation was computed by taking Pearson’s correlation coefficient between the ΔF/F trace and animal’s smoothed velocity with a −5 sec to +5 sec rolling lag. The highest absolute value of the correlation coefficients in this time window was taken as the correlation coefficient for the respective cell.

##### Run-Start/Run-Stop

Based on a velocity cut-off of 0.2 cm/s, run-start and run-stop events were observed throughout the imaging session. Each run-start and run-stop event was at least 10 imaging frames (1-2 seconds) and nearby events within 20 frames (3-5 seconds) were merged. These responses were normalized using a z-score over the interval for each individual cell. Peri-event time averages were generated for each animal. The response magnitude for each cell was computed by taking the difference between the mean post-event ΔF/F and the mean pre-event ΔF/F.

##### Peri-Event Time Averages (PETA)

For salient stimuli such as run-start/run-stop events and reward distribution events, a PETA was constructed around each event respectively. These events were z-scored and then averaged for each respective cell, and the z-scored averages were visualized using a heatmap or an averaged line-plot. A Post-Pre metric was calculated based on the z-scored averages as a measure for event modulation.

##### Position decoding

A linear classifier support vector machine (SVM) was trained on the tuning curves of each respective cell, and the decoding error was visualized. The SVM was cross-validated by training the classifier on (n-1) laps and evaluating on the held-out lap, and this process was repeated such that every lap was utilized as a test lap at least once. For these initial comparisons, all of cells and laps were utilized, which were shown to not be significantly different between the two conditions. To normalize comparisons between conditions, the SVM was bootstrapped with different matched numbers of input cells and input laps averaged by animal to compare the position decoding efficacy between genotypes.

##### Spatial tuning curve

The 2-meter belt was divided into 100 bins (∼2cm per bin). Spatial tuning curves for each cell during an experiment were generated by averaging the velocity regressed ΔF/F (as described above in preprocessing of Ca2+ imaging data) in each bin when the animal was running over 5 cm/s.

##### Peak place field calculation

The spatial tuning curves were shuffled 1000 times by circularly rotating the velocity-regressed ΔF/F tuning curves around the 100 spatial bins to generate a shuffle distribution. Cells were classified as being spatially selective (i.e., having a place field) if the cell’s tuning curve exceeded the 95th percentile (or was under the 5th percentile for negatively selective cells) with at least 10 sequential bins over/under the cutoffs from the shuffled distribution. The number of bins over or under the 95th and 5th percentile curves were visualized and a cutoff of 30 consecutive bins was used for the figures as it represented the approximate 75th quartile for place field width.

##### Spatial information

Using the spatial tuning curve, a spatial information score was computed following the methodology described by Skaggs et al 1992^118^. While the method was originally conceived for spiking data, we applied it onto the spatial tuning curve from Ca^2+^ data as a proxy for information content. As the Skaggs information method utilizes a firing rate tuning curve binned on location, we applied the formula to each interneuron’s respective tuning curve binned on location:

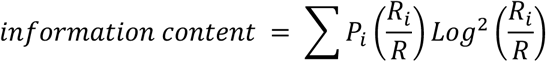

where *i* is the bin number, *P*_i_ is the probability for occupancy of bin *i*, *R*_i_ is the tuning curve for bin *i*, and *R* is the overall mean tuning curve.

##### Reward Modulated Activity

For goal-directed experiments, the peri-event time averages were constructed around each reward event. These responses were normalized using a z-score over the interval for each individual cell. The 2-meter belt was divided into 100 bins (∼2cm per bin). Tuning curves were generated by averaging the ΔF/F in each bin for both regular ΔF/F and velocity-regressed 11F/F.

##### Reward Modulation Index

A reward modulation index (RMI) was computed as described in refs. ^97,98^. Briefly, we compared the difference of the average activity 10 cm preceding the reward zone and the average activity on the remainder of the belt, normalized by the sum of the average activity over the two segments.

##### Pairwise Pearson’s correlation analysis

Pairwise Pearson’s correlation coefficients were computed for every pair of cells between a given cell type pair within a given session. The average coefficient value for each subtype was then calculated per session pair and averaged across sessions within condition. The microcircuit level analysis was averaged across mice. For each matrix when visualized, comparisons were done with the cell type on the left to the cell types on the bottom, as we investigate how each cell type relate to other cell types between animals with differing numbers of cells per type. All directional statistics are included in Supplementary Table 3.

##### Multivariate ridge regression analysis

A multivariate ridge regression model was trained on the ΔF/F trace of Ca^2+^ activity (velocity not regressed) for each cell with three predictors: velocity, position, and licking. Each predictor was normalized to a 0-1 scale by using a minimum-maximum scaler. Given the inherent relationship between these variables, we used a ridge regression model with a cross-validated alpha value to limit the interaction between the predictors.

#### Probabilistic Graphical Model Framework

##### Preprocessing with NMF to remove shared variance

We used non-negative matrix factorization (NMF) here for preprocessing to eliminate sources of common structure to all traces in an unbiased way. For all ΔF/F_0_ traces for each session, we learn a rank-K estimation of the log-transformed signal and then subtract this out. Log transformation is to estimate a multiplicative correction rather than an additive one. We assume that the first K-components that are shared across all cells do not reflect specific cell-to-cell interactions but rather common input patterns and other confounds such as velocity and movement. In mathematical terminology, if we term 𝑓_!,$_ as the ΔF/F_0_ value for the i^th^ cell at the t-^th^ time step, we model the signal as a realization of the following model

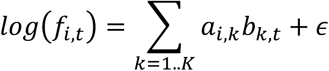

After estimating, **a,b** for each session, we compute the residual and estimate the background subtracted ΔF/F_0_ , 𝑓^;^_*,$_ as:

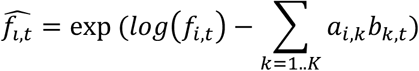

In our experiments, we found K=5 to be a sufficient rank to remove the background effects that are due to factors including but not limited to velocity, mouse head motion, licking and other undefined motor activity. See supplemental figure 9B to see the differences between 𝑓_!,$_ and 𝑓^;^_*,$_. This background subtraction is critical for co-activity analysis since we are looking for subtle hints of direct interactions between cells. For example, if all cells were subjected to global motor pattern signals, this would inflate the baseline correlations between all cells. Thus computing background subtracted signal and using maximum of absolute correlations (used in the Fig. 5 analysis) are two approaches we devised to tease out subtle signals of potentially direct interactions between cells.

##### Model design

We designed a probabilistic graphical model to evaluate functional interactions between neurons, represented by the absolute co-activation between any two cells *i* and *j*, denoted as *x_i,j_*. Each cell *i* was associated with a cell type *c_i_,* drawn from *K* distinct categories. The prior probability of cell *i* belonging to cell type *k* was represented by *p(c_i_ = k) = π_k_*. For each experimental session, the recorded cells were randomly partitioned into two subsets: 80% were allocated to the training set and the remaining 20% to the test set. This split was performed to develop and assess the predictive model in a controlled manner. The labels for the test cells were concealed during the model training phase and were only revealed during the validation stage to verify the accuracy of the model’s identification predictions.

##### Parameterization of cell interactions (cell-to-cell-type co-activity term)

Interactions between each cell and others of a given cell type *k* were modeled using the order-statistics of the empirical distribution of absolute correlations, specifically the maximum. Namely, if 𝑓̂_*i*_ is the background corrected trace of cell i, and 𝑓̂_*i*_ is the background corrected trace of cell j with 𝑐_*i*_, denoting its cell-type, we quantify the interaction of cell i with cell type k as

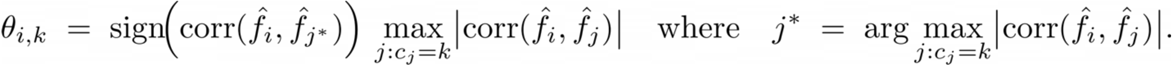

For example, if there is at least one cell from type *k* is highly correlated with cell i, the maximum correlation between between cell i to cell type k is going to be high. Using the approach in fig 4 where average correlation is calculated, this signal will be not captured. Furthermore, we look for unsigned correlations. For example, if cell i has correlation of −0.9 with a cell of type k, and has 0.3 correlation with remainder of cells from type k, our θ_-_, k statistic will be −0.9. This parameterization is chosen to reflect subtle but strong interactions between cell *i* and cell type *k*, capturing the direction and intensity of these inter-cell-type interactions and modeling the interactions as a random distribution to make the model fully Bayesian. Comparisons are directional, but for simplified visualization, a single weighted line is used based on the arithmetic average between the two cell types. Directional arrows are included to illustrate significant effects between cell types, but all directional statistics are included in Supplementary Table 4.

##### Multivariate distribution of interaction parameters (cell-type-to-cell-type co-activity term)

The full interaction profile of cell *i* across all *K* cell types was modeled using a multivariate Gaussian distribution, conditional on the cell type *c_i_*. This distribution is defined by a mean vector *µ_a_* and a covariance matrix Σ_a_, specific to cell type *a*, and is denoted as *p(θ_i,1_,…,θ_i,K_|c_i_ = a) ∼ N(Ω_a_)*. This approach captures the ensemble of interactions that a cell of type a typically engages in with the spectrum of other cell types within the circuit.

##### Inference and Bayesian approach

We infer model parameters using Bayesian inference and computing the maximum a posteriori estimates of cell type *c_i_* for a subset of cells *{c_j_}_j∈L_*, where *L* is the set of indices corresponding to known cell types. This process leveraged both the prior probabilities encoded in the model and the observed cell-to-cell interactions to probabilistically assign cell types to the remaining cells in the circuit. These analyses were performed for *Df(16)A^+/-^* and WT mice, separately.

#### Neuronal identity prediction

##### Classification validation

Our model was validated through its ability to classify cell identity based on functional correlation patterns. Model validation was performed by comparing the predictive accuracy of our probabilistic model against a null distribution. The null distribution was generated by shuffling cell type labels, thereby preserving the overall structure of the data while eliminating any true correlation between cell types and their functional interaction patterns. This approach allowed us to control for the possibility that any predictive success of the model could be due to chance. Predictive accuracies of our model were then contrasted with those from the null distribution to affirm the model’s capability to identify cell types beyond random assignment. To assess the robustness of our probabilistic model, we conducted 31 independent runs, each initialized with a different random seed.

### QUANTIFICATION AND STATISTICAL ANALYSIS

#### Statistical analyses

Subtype comparisons were conducted using one-way ANOVA tests with Tukey’s range tests with correction for multiple testing if appropriate. For comparisons across genotypes, unpaired t tests were utilized if the data followed a normal distribution, assessed using the Shapiro-Wilk test. Otherwise, for non-normally distributed comparisons, the Mann-Whitney U test was used. The Kolmogorov–Smirnov test was used when comparing CDFs of continuous data. For pairwise correlation analysis, we performed Bonferroni correction for the number of comparisons performed (multiplying the computed p-values by 25). For comparisons that had to preserve multiple features, we performed a Hotelling’s T-squared distribution test. P < 0.05, **: P < 0.01, ***: P < 0.001. n.s. is not significant. All data analysis and visualization were done with custom software built on Python version 2.7.15, 3.8.11, and MATLAB R2023b.

## Acknowledgements

S.A.H. is supported by the Burroughs Wellcome Fund and the Stavros Niarchos Foundation. A.L. is supported by NIMHR01MH124047, NIMHR01MH124867, NINDSR01NS121106, NINDSU01NS115530, NINDSR01NS133381, NINDSR01NS131728, NIARF1AG080818. E.V. is supported by NIMHR00MH128772. J.A.G. is supported by NIMHR01MH124047, NIMH R01MH097879 and the Stavros Niarchos Foundation. We thank Tristan Geiller, Bert Vancura for consultation on the analysis, and Erica Rodriguez for the mouse schematic in Fig. 1A.

## Author Contributions

S.H., B.R., J.A.G., and A.L. conceived the study and wrote the manuscript. S.H. and B.R. performed experiments and analyzed the data with assistance from M.E.C.P., A.T., and H.A.. S.H. and E.V. conceived the coactivity and CATT analyses, which was developed and implemented by E.V., M.E.C.P., and H.A..

---

**Supplementary Table 1:**
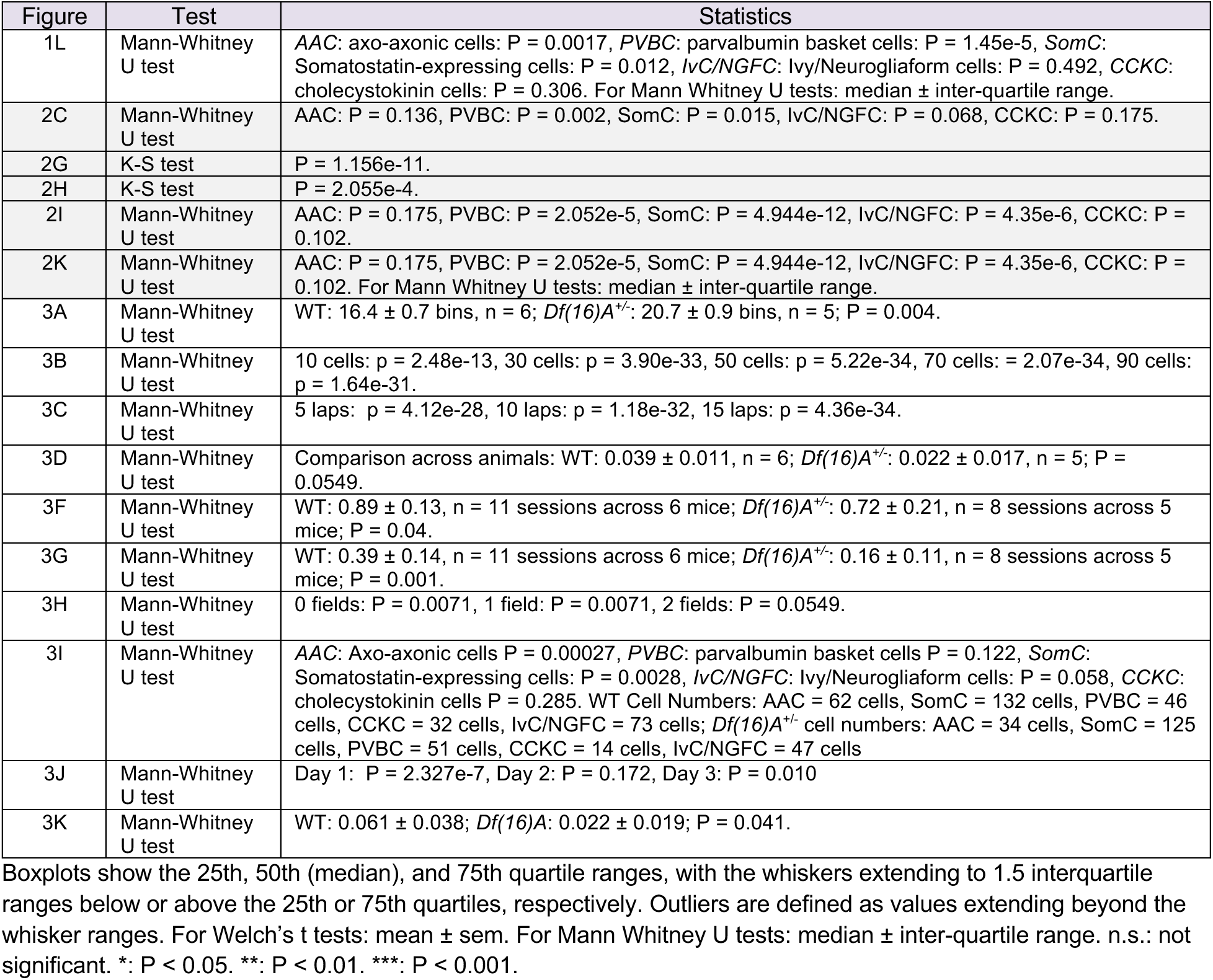
Statistical detail summary.

**Supplemental Figure 1.**
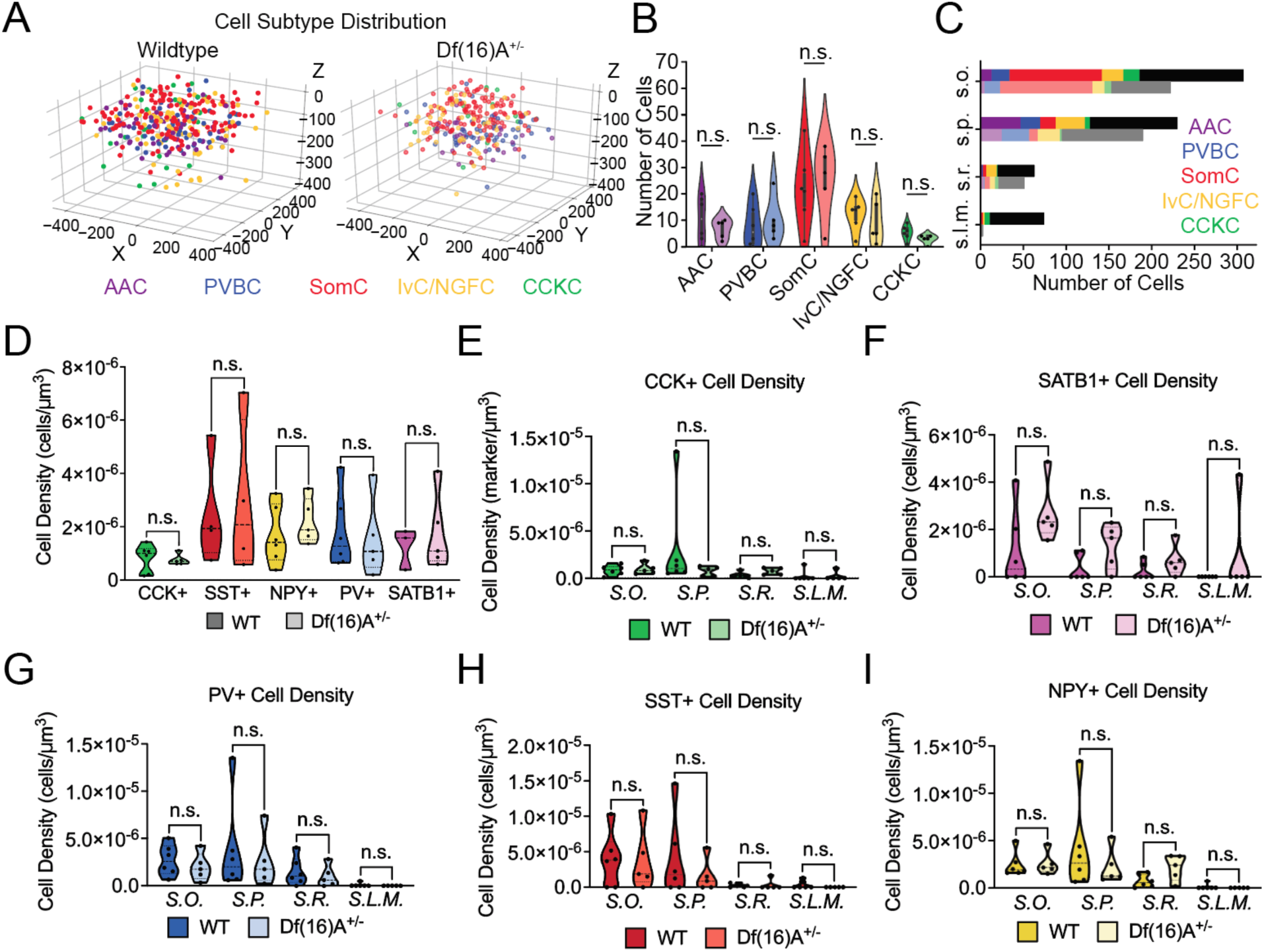
3D-AOD imaging and molecular characterization strategy for interneuron subtypes. **A.** Summary distribution plots of all cells imaged in this study for each genotype based on imaging coordinates superimposed across all animals (n = 6 WT, n = 5 *Df(16)A^+/-^* mice) Dimensions are in microns (800 µm x 800 µm x 400 µm). **B.** Cell type number totals averaged per genotype (n = 6 WT, *n* = 5 *Df(16)A^+/-^* mice) (Mann-Whitney U Test, p > 0.05). **C.** Distribution of total cells identified, ordered by cell type, across all layers of CA1. *s.l.m.: stratum lacunosum moleculare; s.r.: stratum radiatum; s.p.: stratum pyramidale; s.o.: stratum oriens.* Black and gray bars represent cells that could not be identified **D.** Quantification of total marker density for all immunohistochemical markers used (unpaired t-tests with Welch’s correction (P > 0.05)). **E.** Quantification of the density of cholecystokinin+ cells within each layer of CA1. *CCK*: Cholecystokinin. 6-7 serial 75 μm sections were stained and quantified for CCK+/GCaMP+ cell signal across the four layers of CA1 in n = 6 WT n = 6 *Df(16)A^+/-^*mice (unpaired t-tests with Welch’s correction (P > 0.05)). **F.** Quantification of the density of SATB1+ cells within each layer of CA1. 6-7 serial 75 μm sections were stained and quantified for SATB1+/GCaMP+ cell signal across the four layers of CA1 in n = 6 WT, n = 6 *Df(16)A^+/-^*mice (unpaired t-tests with Welch’s correction (P > 0.05)). **G.** Quantification of the density of parvalbumin+ cells within each layer of CA1. *PV*: parvalbumin. 6-7 serial 75 μm sections were stained and quantified for PV+/GCaMP+ cell signal across the four layers of CA1 in n = 6 WT, n = 6 *Df(16)A^+/-^* mice (unpaired t-tests with Welch’s correction (P > 0.05)). **H.** Quantification of the density of somatostatin+ (*Som*) cells within each layer of CA1. 6-7 serial 75 μm sections were stained and quantified for Som+/GCaMP+ cell signal across the four layers of CA1 in n = 6 WT n = 5 *Df(16)A^+/-^*mice (unpaired t-tests with Welch’s correction (P > 0.05)). **I.** Quantification of the density of NPY+ cells within each layer of CA1. 6-7 serial 75 μm sections were stained and quantified NPY+/GCaMP+ cell signal across the four layers of CA1 in n = 6 WT, n = 6 *Df(16)A^+/-^* mice (unpaired t-tests with Welch’s correction (P > 0.05)). *NPY*: Neuropeptide Y. *n.s.*= not significant.

**Supplemental Figure 2.**
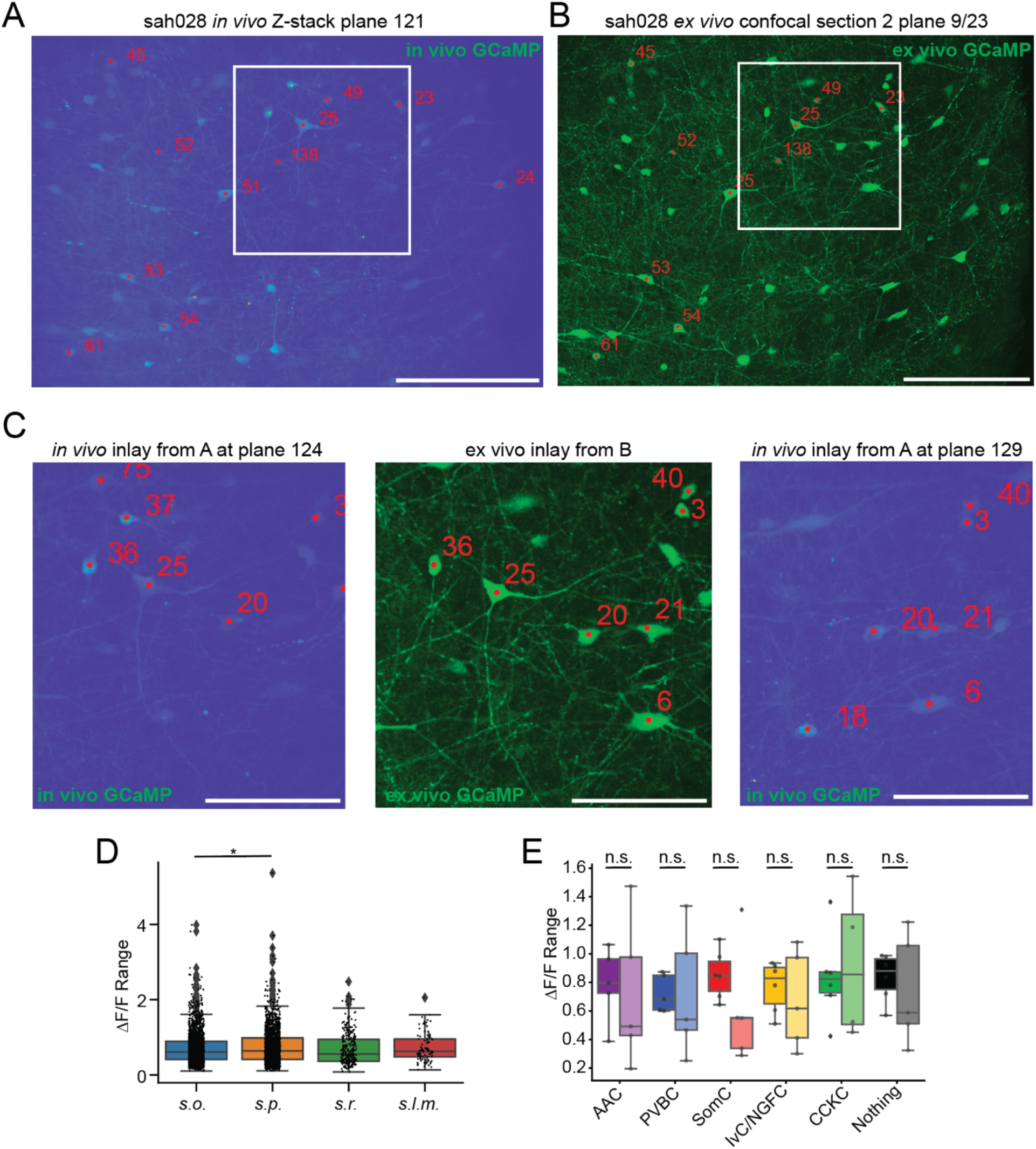
Example detailed registration and imaging. **A.** Plane 121 out of 135 from the coordinate labeled *in vivo* z-stack for mouse sah028. Scale bar is 200 µM. Numbered cells denote coordinate locations in X,Y,Z taken from the *in vivo* imaging ROIs and mapped onto the Z-stack. **B.** *Ex vivo* 75 µM confocal section #2 from mouse sah028 transverse sectioning. Transverse sections are taken to match the imaging plane taken *in vivo*. Numbered cells denote manually matched cells imaged *in vivo*. Scale bar is 200 µM. **C.** Middle panel represents the inlay from the confocal section depicted in B. Scale bars are 100 µM. Left and Right panels depict moving in Z in the Z-stack to match imaged cells in the section to plane 124 (left) and plane 129 (right). **D.** One-way ANOVA (p = 0.007) with post hoc Tukey’s range test corrected for multiple testing. Cells in the s.p. had greater dynamic range than s.o. (p = 0.0054.), all other comparisons were n.s.. **E.** dF/F range between genotypes by cell type across animals (Mann-Whitney U test: SomC: P = 0.06, AAC: P = 0.42, PVBC: P = 0.34, IvC/NGFC: P = 0.46, CCKC: P = 0.46, Nothing: P = 0.39). s.o.: stratum oriens; s.p.: stratum pyramidale; s.r.: stratum radiatum; s.l.m.: stratum lacunosum moleculare. n.s.: not significant. *: P < 0.05.

**Supplemental Figure 3.**
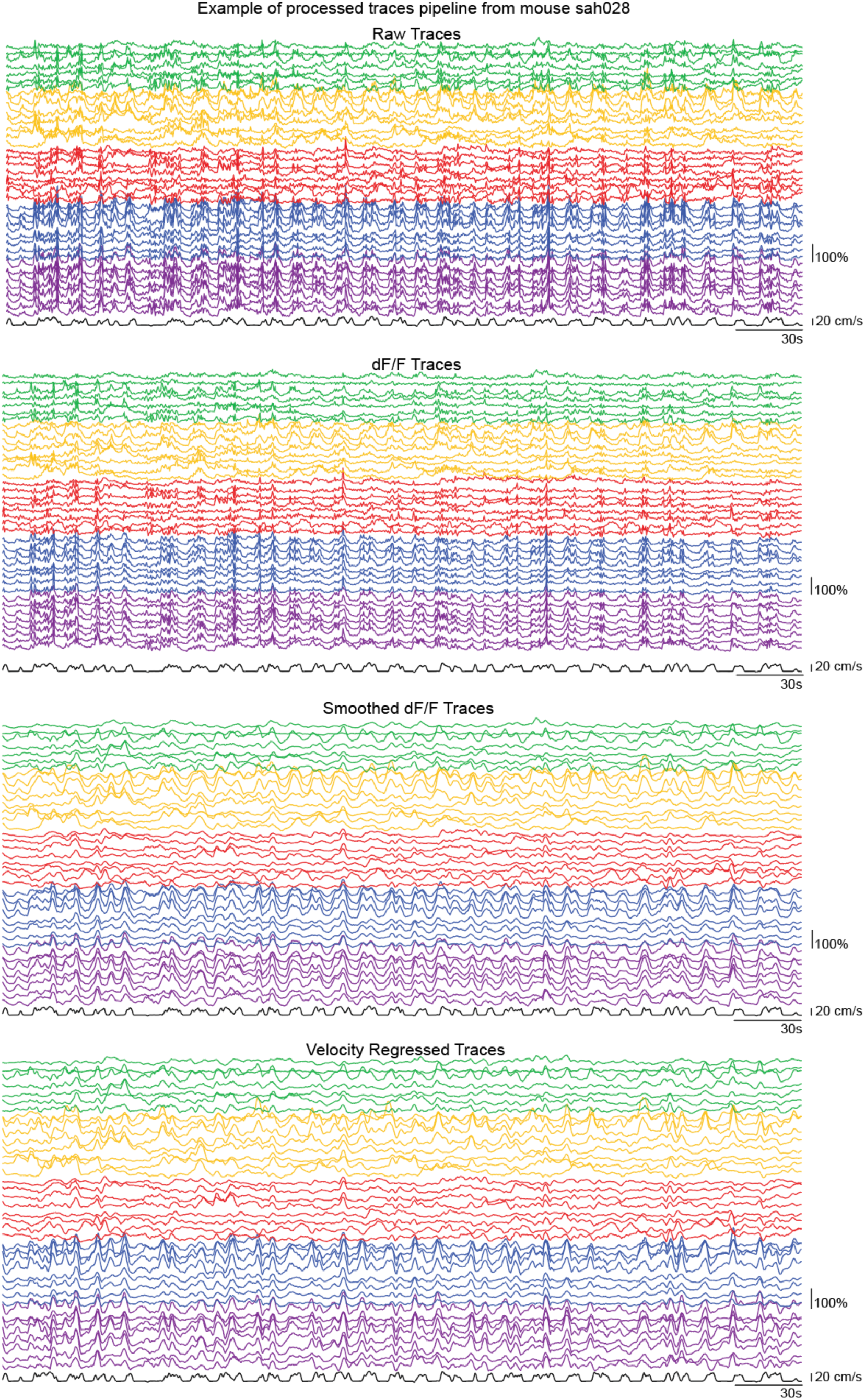
Example processed traces for analysis for. **figures 1-4**. Depicted here are 8 example traces per cell type from the animal number sah028. ΔF/F was computed based on a 1st percentile baseline value over a 30-second rolling window on the raw traces extracted from our manually curated regions of interest (ROIs). The ΔF/F traces were then smoothed using a Savitzky-Golay filter with a window length of 9 and a polyorder of 3. For analysis involving velocity-regressed traces, each animal’s velocity was smoothed with a Hanning filter with a window of 10, and then covariate linear regression was applied to each fluorescence trace where the smoothed velocity was regressed out.

**Supplemental Figure 4.**
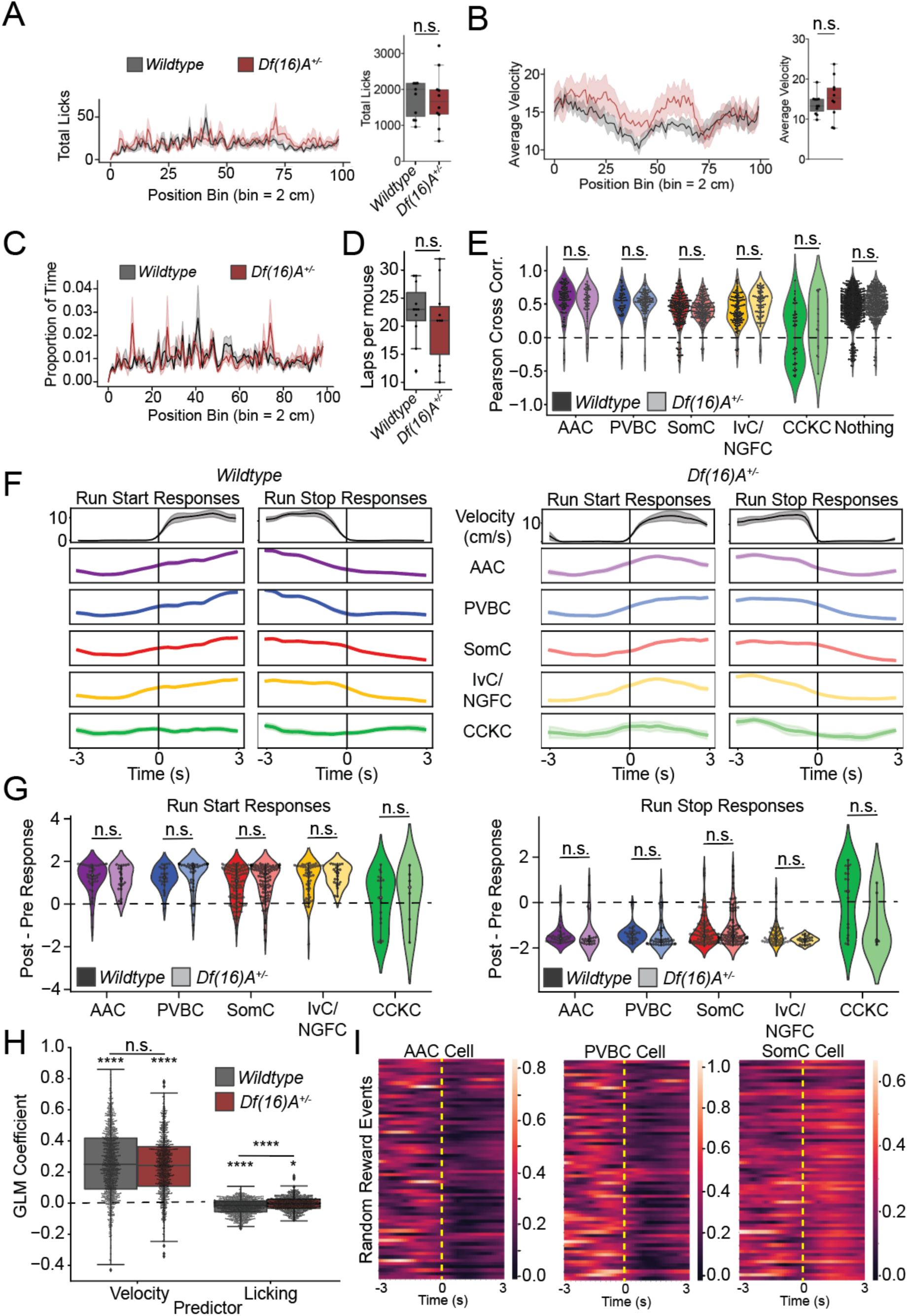
Locomotion related interneuron activity is preserved in Df(16)A^+/-^ mice. **A.** Licking proportion of WT (black) and *Df(16)A^+/-^* (brown). There is no significant difference in lick location (left) or total licks (right) across the belt during Random Foraging in WT (*n* = 6) or *Df(16)A^+/-^* (*n* = 5) depicted by box-and-whisker plot, WT: 1534 ± 190 licks, *Df(16)A^+/-^*: 1608 ± 291 licks, Welch’s t-test, p = 0.838. B) Average velocity in cm per second plotted by location on the belt of WT (black) and *Df(16)A^+/-^* (brown) mice (left). There is no significant difference in average velocity during Random Foraging in WT (n = 6) or *Df(16)A^+/-^* (n = 5) depicted by box-and-whisker plot (right), WT: 13.7 ± 0.7 cm/s, *Df(16)A^+/-^*: 15.2 ± 1.6 cm/s, Welch’s t-test, P = 0.405. **C.** Total laps across mice within genotype (WT: 22.6 ± 1.5 laps, *Df(16)A^+/-^*: 20.6 ± 2.2 laps, Welch’s t test, t = 0.74, P = 0.47). **D.** Average percent occupancy on the belt by position per genotype (Mann-Whitney U test, P = 0.3). **E.** Pearson correlation coefficients of interneuron activity and velocity for all tested interneuron subtypes (WT: n = 97 AACs, 53 PVBCs, 194 SomCs, 115 IvCs/NGFCs, and 44 CCKCs from n = 6 mice, *Df(16)A^+/-^*: 46 AACs, 87 PVBCs, 181 SomCs, 72 IvCs/NGFCs, and 14 CCKCs from n = 5 mice). WT: one-way ANOVA (p < 10^26^), *Df(16)A^+/-^*: one-way ANOVA (P < 10^10^) with post hoc Tukey’s range test corrected for multiple testing. **F.** Peri-event time averages (PETAs) of averaged z-scored traces with a range between −1.75 and 1.75 z-scores during Run-Start and Run-Stop events. Left: WT; Right: *Df(16)A^+/-^*. **G.** Average Post-Pre PETA responses to Run-Start (Left) and Run-Stop (Right) events. **H.** Ridge Regression Model shows velocity as a significant predictor of IN GCaMP-Ca^2+^ activity. Wilcoxon: WT: 0.257 ± 0.007, P = 4.68e-141, *Df(16)A^+/-^*: 0.240 ± 0.007, P = 3.60e- 117. **I.** Example per event rasters for individual cell responses to reward onset (yellow dashed line) during the random foraging task. Left: AAC cell; Middle: PVBC cell; Right: SomC cell. Boxplots show the 25th, 50th (median), and 75th quartile ranges, with the whiskers extending to 1.5 interquartile ranges below or above the 25th or 75th quartiles, respectively. Outliers are defined as values extending beyond the whisker ranges. For Welch’s t tests and Wilcoxon test: mean ± sem. For Mann Whitney U tests: median ± inter-quartile range. n.s.: not significant. ****: P < 0.0001.

**Supplemental Figure 5.**
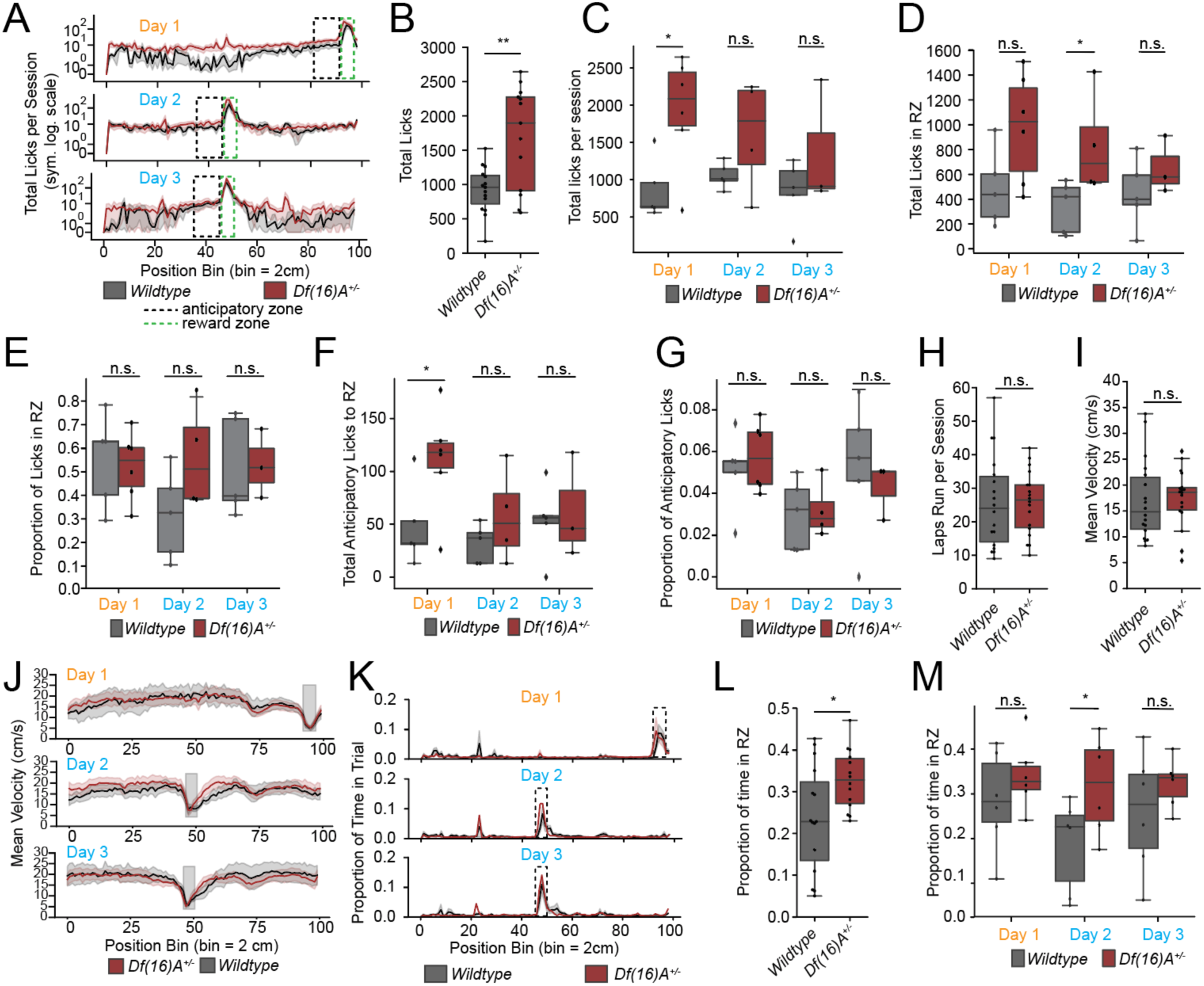
Mutant mice exhibit protracted behaviors without significant alterations in task performance. **A.** Total lick distribution across position bins on familiar session Day 1 (top), translocation Day 2 (middle) and translocation Day 3 (bottom) of recording (symmetrical log scale). **B.** Total licks across all sessions (WT: 959 ± 415 licks; *Df(16)A^+/−^*: 1895 ± 1364 licks; Mann-Whitney U test: P = 0.0064). **C.** Total licks across sessions per day (D1: WT: 861 ± 180 licks; *Df(16)A^+/−^*: 1928 ± 306 licks; Mann-Whitney U test: P = 0.019). (D2: WT: 1054 ± 76 licks; *Df(16)A^+/−^*: 1612 ± 381 licks; Mann-Whitney U test: P = 0.15). (D3: WT: 848 ± 188 licks; *Df(16)A^+/−^*: 1367 ± 487 licks; Mann-Whitney U test: P = 0.28). **D.** Total licks in reward zone (D1: WT: 484 ± 138 licks; *Df(16)A^+/−^*: 976 ± 180 licks; Mann-Whitney U test: P = 0.06). (D2: WT: 341 ± 93 licks; *Df(16)A^+/−^*: 834 ± 210 licks; Mann-Whitney U test: P = 0.03). (D3: WT: 445 ± 124 licks; *Df(16)A^+/−^*: 654 ± 133 licks; Mann-Whitney U test: P = 0.19). **E.** Proportion of licks in reward zone (D1: WT: 0.547 ± 0.088; *Df(16)A^+/−^*: 0.523 ± 0.59; Mann-Whitney U test: P = 0.39). (D2: WT: 0.317 ± 0.84; *Df(16)A^+/−^*: 0.563 ± 0.111; Mann-Whitney U test: P = 0.09). (D3: WT: 0.513 ± 0.092; *Df(16)A^+/−^*: 0.530 ± 0.85; Mann-Whitney U test: P = 0.5). **F.** Total anticipatory licks before the reward zone (D1: WT: 48 ± 17 licks; *Df(16)A^+/−^*: 111 ± 20 licks; Mann-Whitney U test: P = 0.045). (D2: WT: 32 ± 8 licks; *Df(16)A^+/−^*: 56 ± 22 licks; Mann-Whitney U test: P = 0.27). (D3: WT: 53 ± 16 licks; *Df(16)A^+/−^*: 62 ± 29 licks; Mann-Whitney U test: P = 0.76). **G.** Proportion of anticipatory licks (D1: WT: 0.051 ± 0.009; *Df(16)A^+/−^*: 0.057 ± 0.007; Mann-Whitney U test: P = 0.46). (D2: WT: 0.030 ± 0.008; *Df(16)A^+/−^*: 0.032 ± 0.007; Mann-Whitney U test: P = 0.45). (D3: WT: 0.052 ± 0.015; *Df(16)A^+/−^*: 0.043 ± 0.008; Mann-Whitney U test: P = 0.28). **H**. Average laps run by mouse per session averaged across animals (WT: 25.5 ± 16.5 laps; *Df(16)A^+/−^*: 25.5 ± 15.5 laps; Mann-Whitney U test: P = 0.40). **I.** Average velocity run by mouse per session averaged across animals: WT: 14.84 ± 10.01 cm/s; Df(16)A^+/−^: 18.61 ± 4.32 cm/s; Mann-Whitney U test: P = 0.24. **J**. Average velocity by position on the belt for Day 1 (top), Day 2 after translocation of the reward (middle), and Day 3 (bottom). **K.** Proportion of time spent across position bins on familiar session Day 1 (top), translocation Day 2 (middle), and translocation Day 3 (bottom) of recording. **L.** Proportion of time spent in the reward zone during the entire session (averaged across sessions WT: 0.228 ± 0.188; *Df(16)A^+/−^*: 0.328 ± 0.108; Mann-Whitney U test: P = 0.02). **M.** Proportion of time spent in the reward zone during the entire session per day (D1: WT: 0.284 ± 0.046; *Df(16)A^+/−^*: 0.341 ± 0.031; Mann-Whitney U test: P = 0.189, D2: WT: 0.188 ± 0.042; *Df(16)A^+/−^*: 0.318 ± 0.044; Mann-Whitney U test: P = 0.046, D3: WT: 0.259 ± 0.056; *Df(16)A^+/−^*: 0.324 ± 0.022; Mann-Whitney U test: P = 0.235). Boxplots show the 25th, 50th (median), and 75th quartile ranges, with the whiskers extending to 1.5 interquartile ranges below or above the 25th or 75th quartiles, respectively. Outliers are defined as values extending beyond the whisker ranges. For Welch’s t tests: mean ± sem. For Mann Whitney U tests: median ± inter-quartile range. n.s.: not significant. *: P < 0.05. **: P < 0.01. ***: P < 0.001.

**Supplemental Figure 6:**
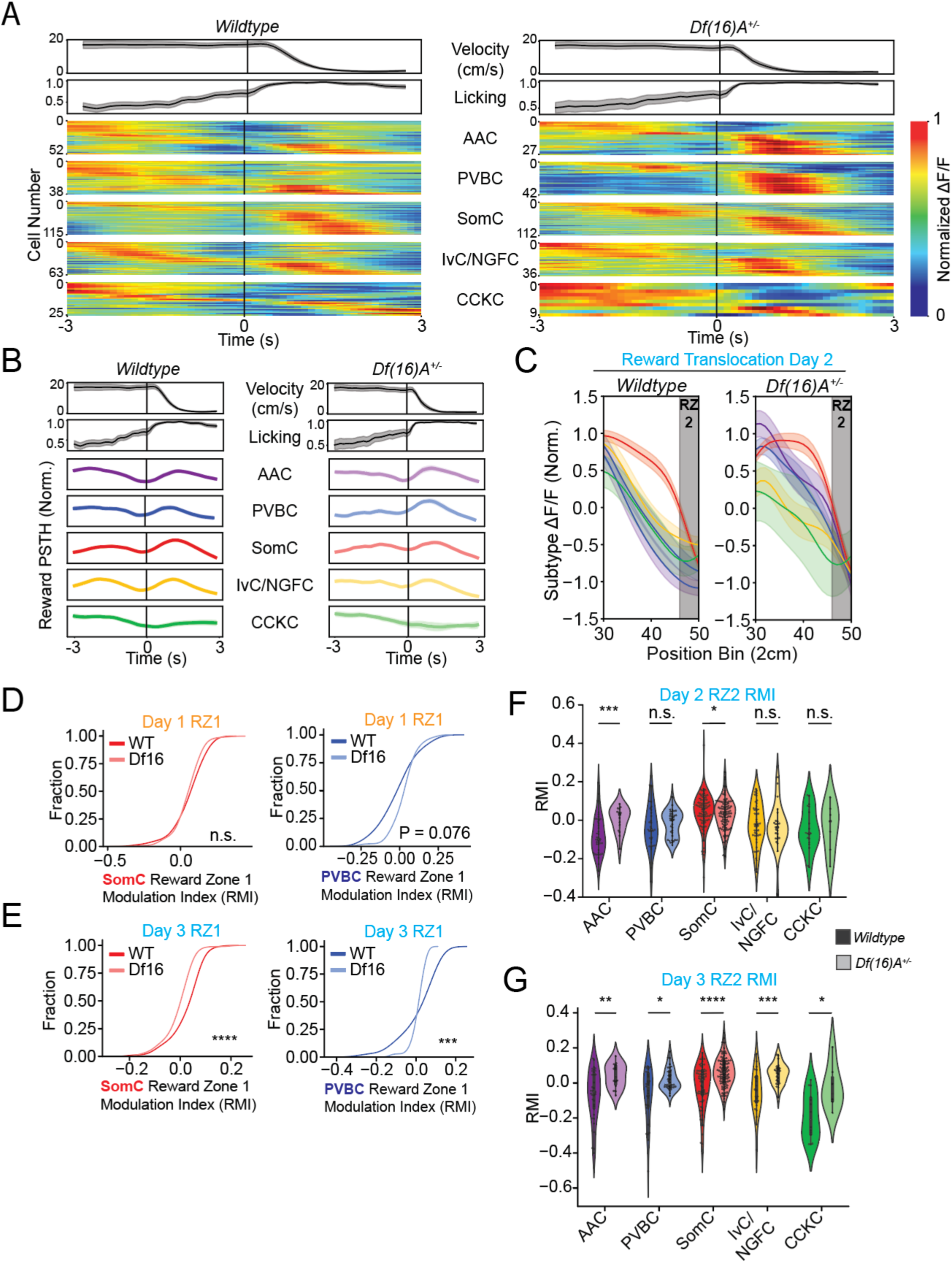
Multiple interneuron subtypes exhibit enhanced reward related activity in mutants. **A.** Individual cell responses to reward during reward translocation task, Left: WT, Right: *Df(16)A^+/-^* **B.** Averaged z-scored cell responses with a range between −1.5 and 1.5 z-scores to reward onset during reward translocation task, Left: WT, Right: *Df(16)A^+/-^*. **C.** Average responses of each cell type in the preceding 40 cm of the belt to the *new* reward location in the WT (left) and *Df(16)A^+/-^* (right) on Day 2. **D.** Reward zone modulation index (*RMI,* see Methods under Reward Modulation Index) for SomCs and PVBCs at the old reward zone during the first day. (SomCs: Kolmogorov-Smirnov (KS) test: P = 0.282, PVBCs: KS test: P = 0.076.) **E.** RMI for SomCs and PVBCs at the old reward zone on second translocation day (Day 3). (SomCs: KS test: P = 3.812e-6, PVBCs: KS test: P = 6.888e-4.) **F.** Reward zone modulation index for all cells at the new reward zone on the first day of reward translocation. (AAC: Mann-Whitney U test: P = 7.742e-4, PVBC: Mann-Whitney U test: P = 0.153, SomC: Mann-Whitney U test: P =0.049, IvC/NGFC: Mann-Whitney U test: P = 0.254, CCKC: Mann-Whitney U test: P = 0.409.) **G.** Reward zone modulation index for all cells at the new reward zone on the second day (Day 3) of reward translocation. (AAC: Mann-Whitney U test: P = 0.0008, PVBC: Mann-Whitney U test: P = 0.153, SomC: Mann-Whitney U test: P = 0.049, IvC/NGFC: Mann-Whitney U test: P = 0.254, CCKC: Mann-Whitney U test: P = 0.409). **F.** Reward zone modulation index for all cells at the new reward zone on the second day (Day 3) of reward translocation. (AAC: Mann-Whitney U test: P = 0.003, PVBC: Mann-Whitney U test: P = 0.018, SomC: Mann-Whitney U test: P =3.4×10^-5, IvC/NGFC: Mann-Whitney U test: P = 0.0005, CCKC: Mann-Whitney U test: P = 0.012) *: P < 0.05. **: P < 0.01. ***: P < 0.001. *n.s.* = not significant.

**Supplemental Figure 7:**
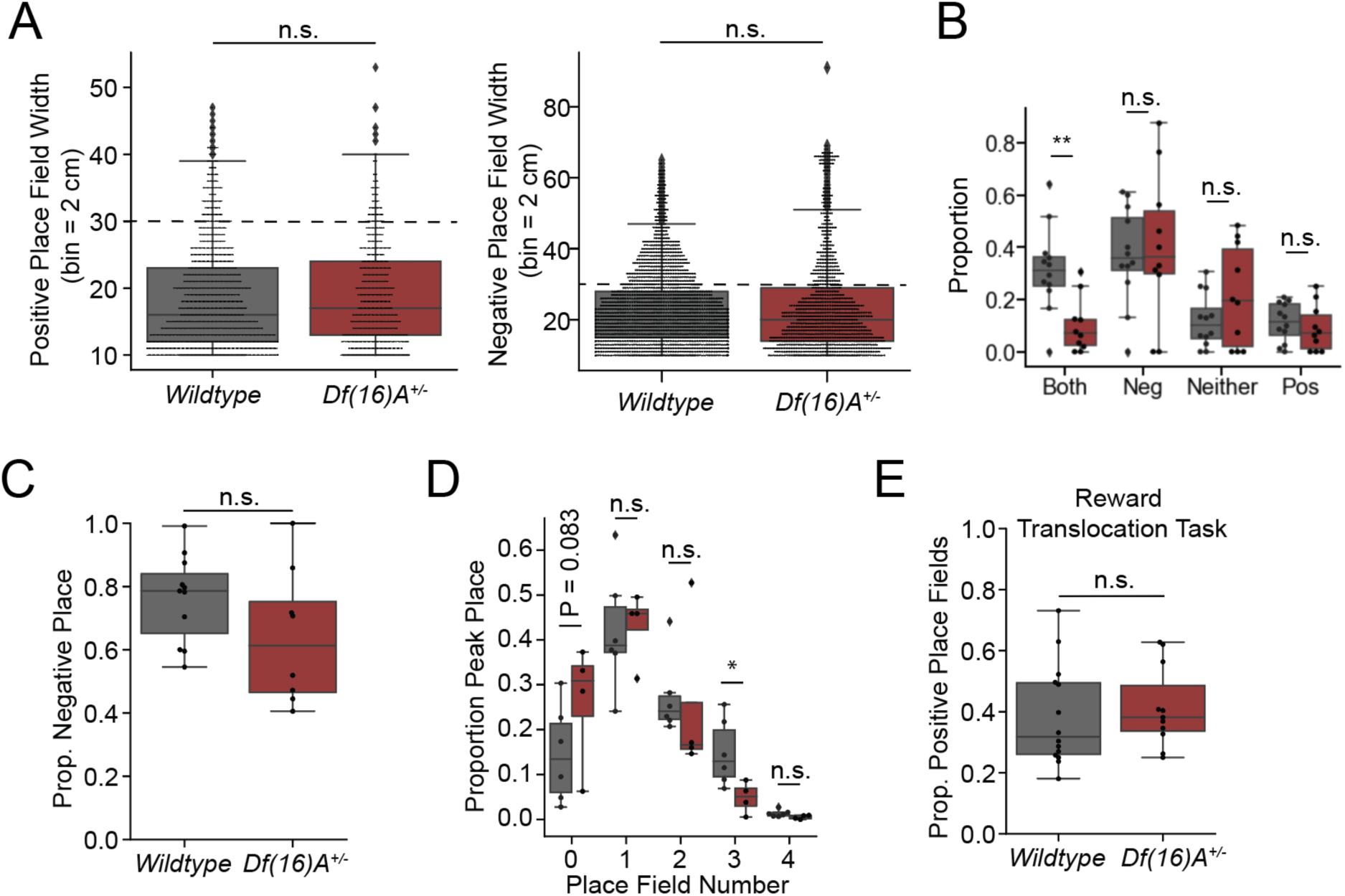
Negative place fields are not reduced in *Df(16)A^+/-^* compared with WT. **A.** Positive (left) and negative (right) place field widths of detected place fields using Peak Place method. Averaged across animals: Mann-Whitney U test: Positive: P = 0.464, Negative: P = 0.375. **B.** Proportions of cells with exclusive types of place fields per mouse, averaged across animals. Mann-Whitney U test: negative (P = 0.63), positive (P = 0.51), both (P = 0.001) and cells with no place fields (0.26). **C.** Negative place fields detected between genotypes. WT: 0.79 ± 0.19, *Df(16)A^+/-^*: 0.61 ± 0.29 Mann-Whitney U test:, P = 0.087. **D.** Number of all detected place fields per cell by genotype. 0 place fields: Mann-Whitney U test: P = 0.0829, 1 place field: Mann-Whitney U test: P = 0.4576, 2 place fields: Mann-Whitney U test: P = 0.1205, 3 place fields: Mann-Whitney U test: P = 0.0126, 4 place fields: Mann-Whitney U test: P = 0.0549. **E.** Proportion of positive place cells during reward translocation task. (Student’s T-test: P = 0.6288). Boxplots show the 25th, 50th (median), and 75th quartile ranges, with the whiskers extending to 1.5 interquartile ranges below or above the 25th or 75th quartiles, respectively. Outliers are defined as values extending beyond the whisker ranges. For Welch’s t tests: mean ± sem. For Mann Whitney U tests: median ± inter-quartile range. *n.s.* = not significant.

**Supplemental Figure 8.**
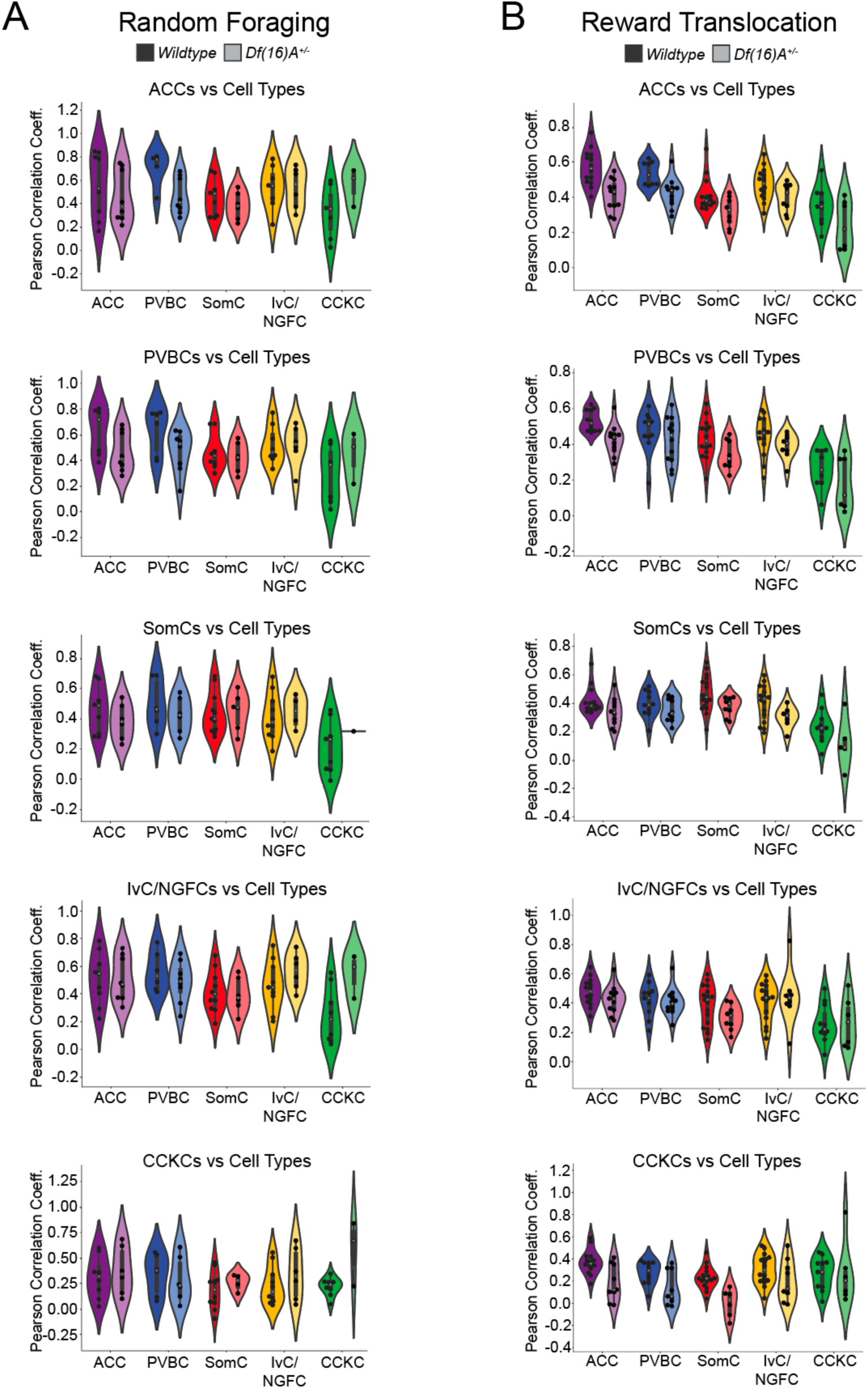
Distribution summaries of correlation coefficients. **A.** Random foraging task sessions pooled. **B.** Reward Translocation task sessions pooled. Detailed information on the inter quartile range of these values can be viewed in Suppl. Table 3.

**Supplemental Figure 9.**
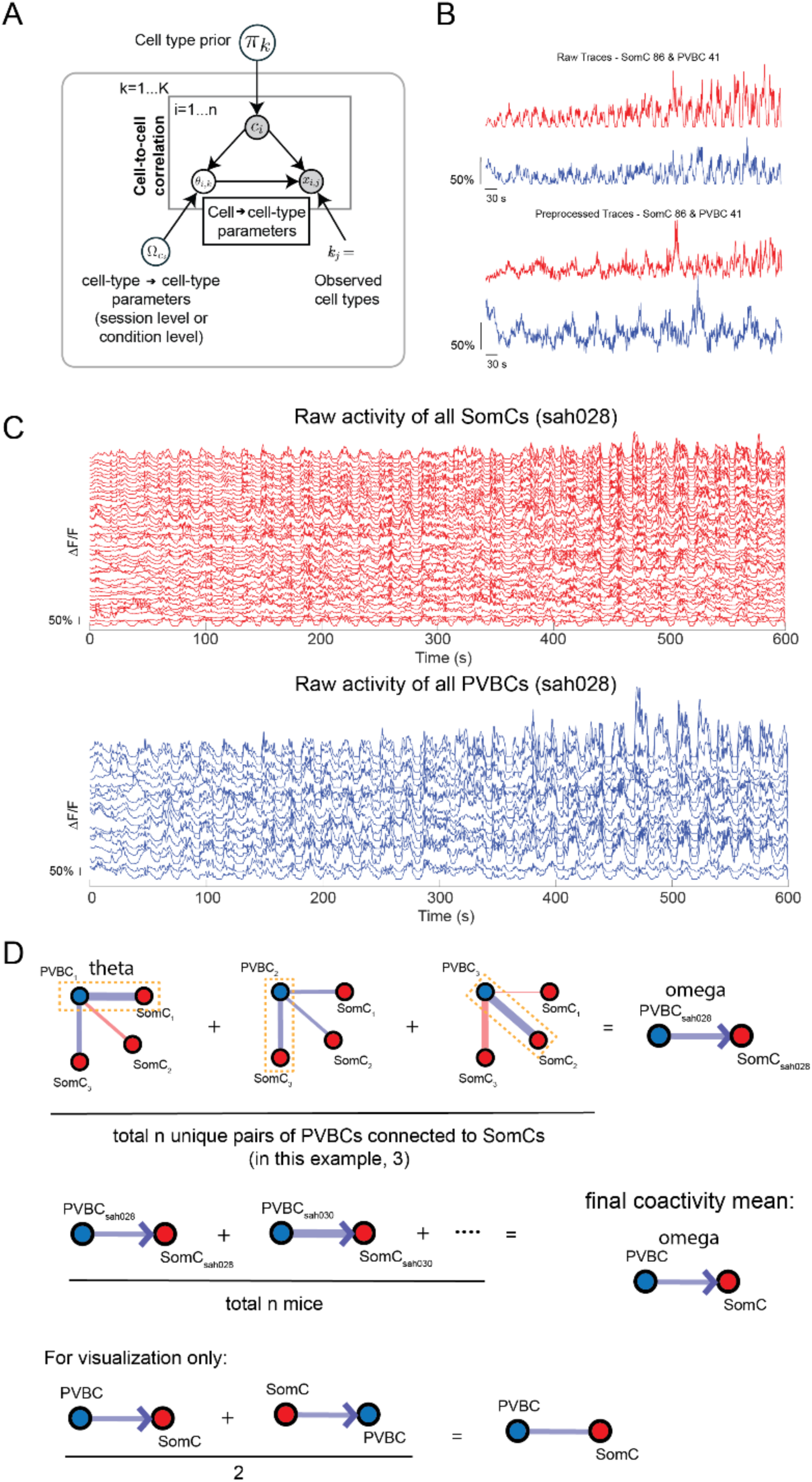
A probabilistic graphical model to infer coactivation patterns. **A.** A schematic of the probabilistic graphical model of cell type to cell type interactions. The model begins with a cell type prior (*π_k_*) for each of the *K* possible types denoting probability of assignment by chance. Each cell (*i*) is then probabilistically assigned a cell type (*c_i_*) using maximum a posteriori estimation, weighed by the prior. Parameters *θ_i,k_* reflect the unique interaction dynamics of cell *i* with cells of type *k* and are derived from cell-to-cell correlations (*x_i,j_*). These parameters are informed by both the observed cell types (*c_ij_* = *k*) and the overarching cell-type-to-cell-type parameters (*Ω_ci_ = [𝜇,*σ]). Arrows depict the direction of dependency of random variables. White circles indicate unobserved random variables, grey circles indicate observed variables (ie, cell type (c_i)_, cell to cell correlation (x_i,j)._ See Probabilistic Graphical Model Framework section for more detail in Methods section. **B.** We process the data, filtering out noise. Cells with variance below a defined threshold (varCutoff) are filtered out, and Non-Negative Matrix Factorization (NMF) removes shared variance and isolates neuron-specific activity. **C.** This pipeline takes raw neural recordings and extracts coactivity features from a pair of cell types within a single session. **D.** We then align and classify neurons and identify the maximum correlation value between a pair of cell types within a given session, and then find the means across animals to build the final structured coactivity graph. For visualization, we average both directions, but all conclusions were made with the direction in mind. The ultimate goal is to extract functional insights from large-scale neural data.

**Supplemental Figure 10.**
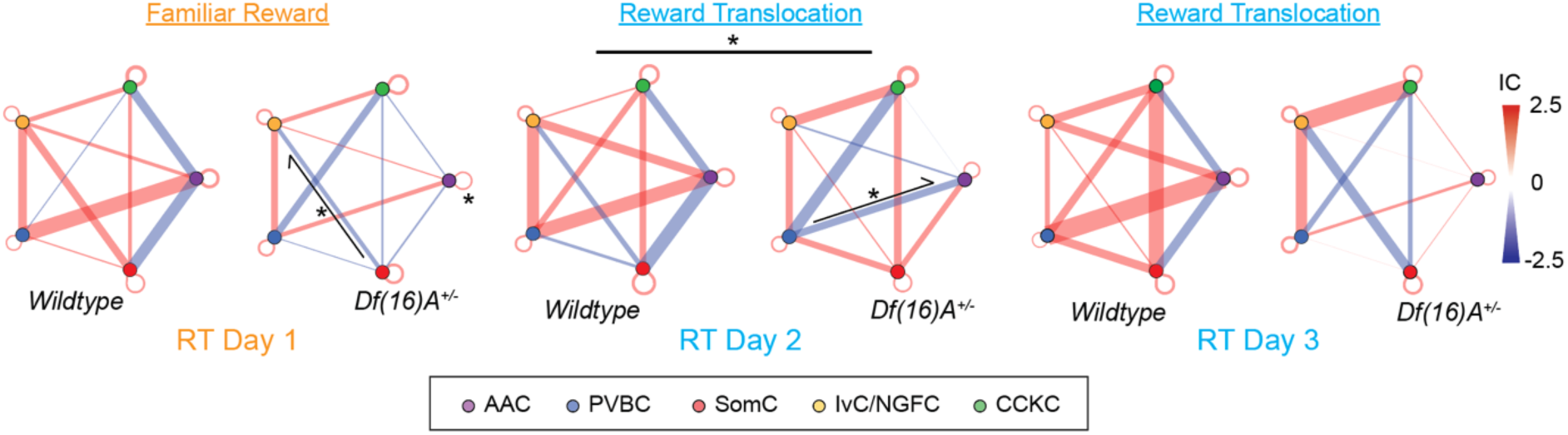
A probabilistic graphical model to infer coactivation patterns. Breakdown of microcircuit and cell type to cell type coactivity structure across days. n.s.: not significant. *: P < 0.05. **: P < 0.01. ***: P < 0.001. See Suppl. Table 4 for detailed statistics.

**Supplemental Figure 11:**
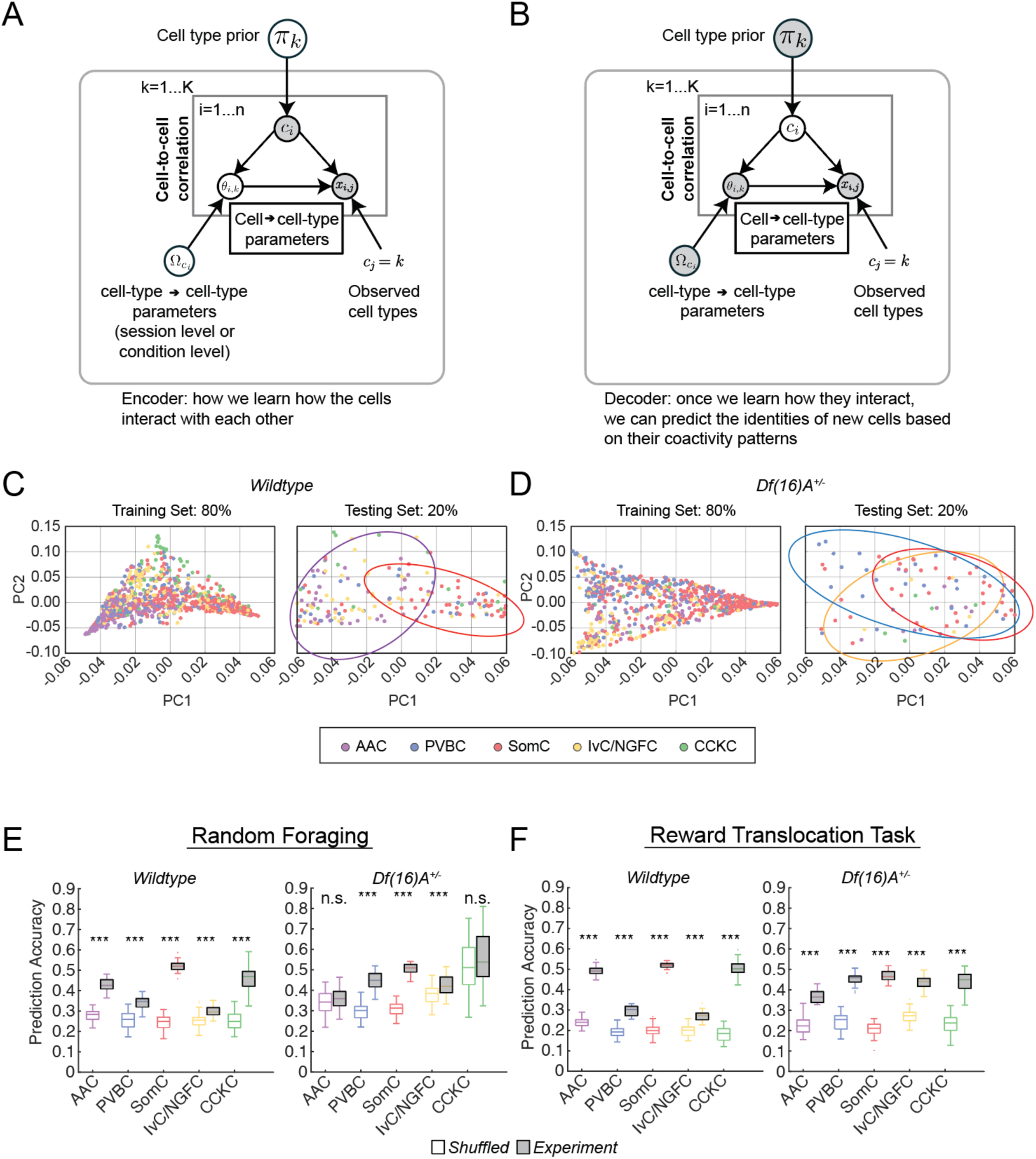
The graphical model describes distinct decoding properties between interneuron subtypes between genotypes. **A.** A schematic of the probabilistic graphical model of cell type to cell type interactions as an encoder. The model begins with a cell type prior (*π_k_*) for each of the *K* possible types denoting probability of assignment by chance. Each cell (*i*) is then probabilistically assigned a cell type (*c_i_*) using maximum a posteriori estimation, weighed by the prior. Parameters *θ_i,k_* reflect the unique interaction dynamics of cell *i* with cells of type *k* and are derived from cell-to-cell correlations (*x_i,j_*). These parameters are informed by both the observed cell types (*c_ij_* = *k*) and the overarching cell-type-to-cell-type parameters (*Ω_ci_ = [𝜇,*σ]). Arrows depict the direction of dependency of random variables. White circles indicate unobserved random variables, grey circles indicate observed variables (ie, cell type (c_i)_, cell to cell correlation (x_i,j)._ See Probabilistic Graphical Model Framework section for more detail in Methods section. **B.** The same schematic, but the connections are known and computed, max coactivity is known, and now we try to estimate cell type (𝑐_𝒾_). **C.** Left: projection of cell-to-cell-type interaction parameters (*θ_i,k_*) for all WT cells with cell types labeled in the training set. Cell types are denoted with the following colors: AAC (purple), CCKC (green), IvC/NGFC (blue), PVBC (yellow), and SomC (red). Right: projection of cell-to-cell-type interaction parameters (*θ_i,k_*) into a low-dimensional subspace for five different hippocampal IN subtypes in the testing set. Large ellipses highlight cell type clusters that appear to drive major principle components. **D.** Same as C but for *Df(16)A^+/-^* cells. (E/F) Cell type classification accuracy using inferred cell types from *A* using our IN activity data for WT and *Df(16)A^+/-^* during **E.** Random Foraging and **F.** reward translocation task. Light boxes indicate the null distribution of classification accuracy upon shuffling cell-type labels. E: P-values of WT for Random Foraging task: AAC (P = 3.48e-27***), PVBC (P = 3.91e-13***), SomC (P = 1.71e-43***), IvC/NGFC (P = 4.27e-07***), CCKC (p = 2.56e-23***). P-values of Df(16)A+/- for Random Foraging task: AAC (P = 3.94e-01, n.s.), PVBC (P = 1.47e-20***), SomC (P = 1.25e-32***), IvC/NGFC (P = 2.27e-04***), CCKC (P = 1.43e-01, n.s.). **F.** Decoding analysis results during the Reward Translocation task. F: P-values of Wildtype for Reward Translocation Task: AAC (P = 8.76e-49***), PVBC (P = 1.58e-23***). SomC (P = 3.46e-53***). IvC/NGFC (P = 8.23e-15***). CCKC (P = 2.16e-40***). P-values of Df(16)A+/- for Reward Translocation Task: AAC (P = 2.52e-22***). PVBC (P = 7.21e-31***). SomC (P = 5.72e-41***). IvC/NGFC (P = 2.66e-29***). CCKC (P = 1.49e-24***).

**Supplemental Figure 12:**
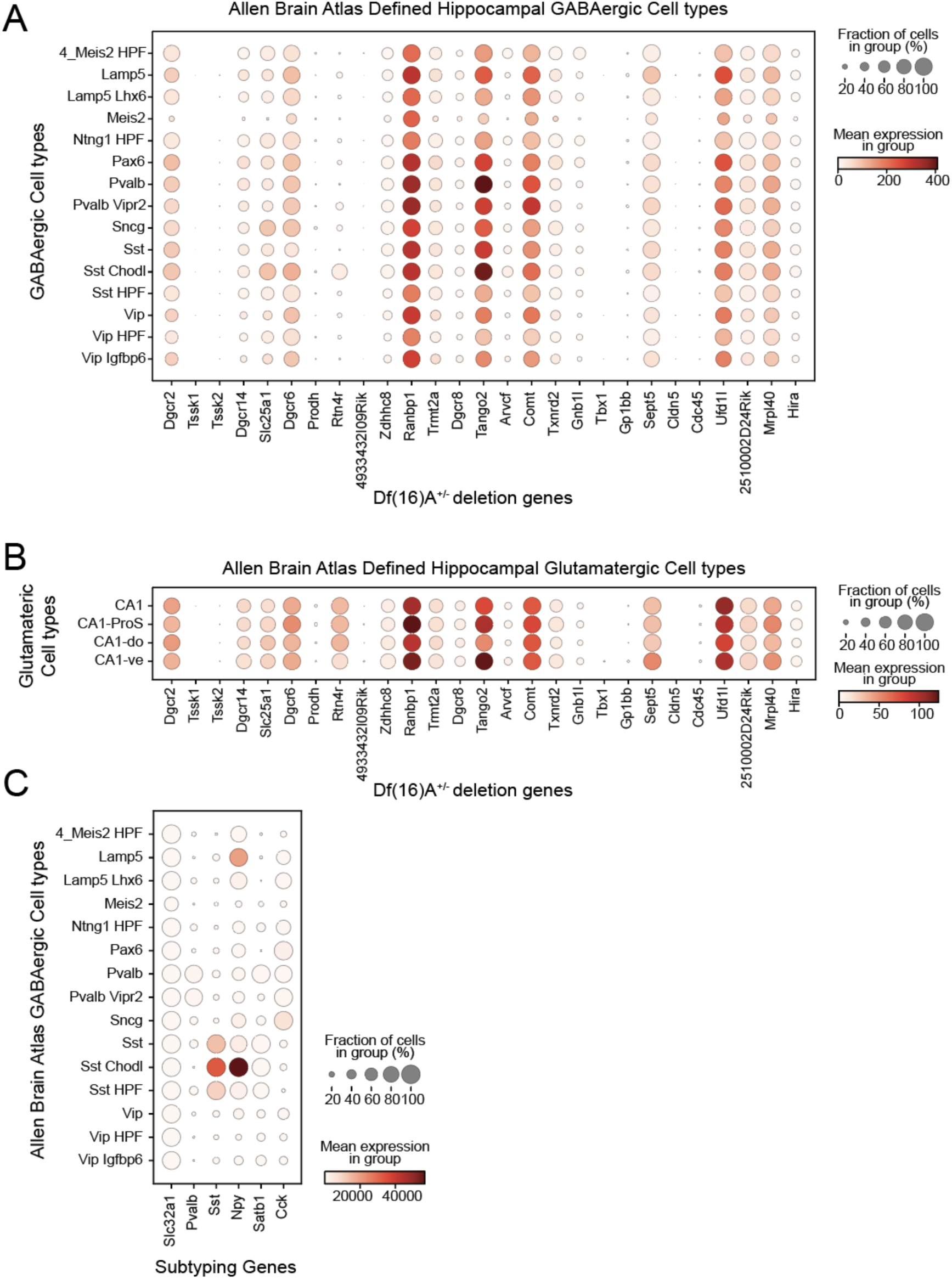
Expression profile of genes residing within the deleted locus in *Df(16)A^+/-^* mice in defined hippocampal. **A.** GABAergic interneuron subtypes and **B.** Glutamatergic subtypes from SMART-seq single cell RNAseq data from the Allen Brain Atlas^159^. **C.** Expression profile of genes used for transgenic and immunohistochemical targeting for cell subtyping in Allen Brain Atlas defined hippocampal GABAergic cell-types.

